# Myristate can be used as a carbon and energy source for the asymbiotic growth of arbuscular mycorrhizal fungi

**DOI:** 10.1101/731489

**Authors:** Yuta Sugiura, Rei Akiyama, Sachiko Tanaka, Koji Yano, Hiromu Kameoka, Shiori Marui, Masanori Saito, Masayoshi Kawaguchi, Kohki Akiyama, Katsuharu Saito

## Abstract

Arbuscular mycorrhizal (AM) fungi, forming symbiotic associations with land plants, are obligate symbionts that cannot complete their natural life cycle without a host. Recently, fatty acid auxotrophy of AM fungi is supported by studies showing that lipids synthesized by the host plants are transferred to the fungi and that the latter lack genes encoding cytosolic fatty acid synthases (1-7). Therefore, to establish an asymbiotic cultivation system for AM fungi, we tried to identify the fatty acids that could promote biomass production. To determine whether AM fungi can grow on medium supplied with fatty acids or lipids under asymbiotic conditions, we tested eight saturated or unsaturated fatty acids (C12–C18) and two β-monoacylglycerols. Only myristate (C14:0) led to an increase in biomass of *Rhizophagus irregularis*, inducing extensive hyphal growth and formation of infection-competent secondary spores. However, such spores were smaller than those generated symbiotically. Furthermore, we demonstrated that *R. irregularis* can take up fatty acids in its branched hyphae and use myristate as a carbon and energy source. Myristate also promoted the growth of *Rhizophagus clarus* and *Gigaspora margarita*. Finally, mixtures of myristate and palmitate accelerated fungal growth and induced a substantial change in fatty acid composition of triacylglycerol compared with single myristate application, although palmitate was not used as a carbon source for cell wall biosynthesis in this culture system. In conclusion, here we demonstrate that myristate boosts asymbiotic growth of AM fungi and can also serve as a carbon and energy source.

**Significance statement:** The origins of arbuscular mycorrhizal (AM) fungi, which form symbiotic associations with land plants, date back over 460 million years ago. During evolution, these fungi acquired an obligate symbiotic lifestyle, and thus depend on their host for essential nutrients. In particular, fatty acids are regarded as crucial nutrients for the survival of AM fungi owing to the absence of genes involved in *de novo* fatty acid biosynthesis in the AM fungal genomes that have been sequenced so far. Here, we show that myristate initiates AM fungal growth under asymbiotic conditions. These findings will advance pure culture of AM fungi.

## Text

Arbuscular mycorrhizal (AM) fungi belonging to the subphylum Glomeromycotina (8) form symbiotic associations with over 70% of land plant species (9). AM fungi provide hosts with minerals taken up via hyphal networks in soil and in return receive carbon sources, such as sugars and lipids derived from plant photosynthates. This is regarded as an obligate symbiotic relationship, as AM fungi can complete their life cycle only through colonization of their host. Nevertheless, a few reports of AM fungal culture without hosts have been published. One report showed that the co-cultivation of the AM fungus *Rhizophagus irregularis* (formerly *Glomus intraradices*) with bacterial strains of *Paenibacillus validus*, separated from each other by dialysis membranes, induced secondary spore formation in the AM fungus (10). Another study reported that some fatty acids, including palmitoleic acid and (*S*)-12-methyltetradecanoic acid (anteiso-C15:0), induced the formation of infection-competent secondary spores in asymbiotic cultures of AM fungi (11). These results suggest that AM fungi may be cultured independently from host plants under artificial conditions. In nature, the life cycle of AM fungi proceeds as follows. Resting spores of AM fungi germinate, and then germ tubes emerge from the spores and elongate into the soil. After colonization into plant roots, AM fungi form highly branched hyphal structures, called arbuscules, in plant cortical cells, which are the sites of nutrient exchange between AM fungi and their hosts. After receiving carbon sources from their hosts, AM fungi activate the formation of extraradical hyphal networks in soil and form spores on their hyphae. Altogether, obtaining carbon sources from their hosts for the production of energy and the carbon skeleton of fungal cell components is a key step for AM fungal growth and reproduction. In particular, tracing experiments and nuclear magnetic resonance (NMR) analyses showed that hexoses are transferred to AM fungi as a carbon source (12-14). Moreover, several AM fungal monosaccharide transporters have been identified (15-17). Recently, lipids have been proven to be another plant-derived carbon source (1-3, 18, 19). Since no fatty acid biosynthesis in extraradical hyphae has ever been detected, lipids were assumed to be synthesized in intraradical hyphae and transferred to extraradical hyphae and spores (14, 20). However, AM fungal genomes do not possess genes encoding cytosolic fatty acid synthases involved in *de novo* fatty acid biosynthesis, indicating that AM fungi cannot produce long-chain fatty acids by themselves (4-7). On the other hand, during AM symbiosis, plants activate fatty acid biosynthesis and transfer lipids, presumably 16:0 β-monoacylglycerol (β-MAG), to AM fungi via arbuscules (1-3, 18, 19). In fact, in plant mutants defective in fatty acid biosynthetic genes that are specific to the pathway supplying lipids to their symbionts, AM fungi cannot develop arbuscules and their root colonization is reduced. These results indicated that AM fungi require exogenous fatty acids for their growth. Once AM fungi take up fatty acids, they can utilize them through fatty acid desaturases and elongases encoded by genes present in their genomes and expressed in intraradical hyphae (4, 5, 7, 20-23). Thus, we examined whether AM fungi can grow and produce fertile spores under asymbiotic conditions through the application of fatty acids.

## Results

### Myristate activates AM fungal growth

Initially, we screened several fatty acids and β-MAGs to identify chemicals that can promote the growth of *R. irregularis* under asymbiotic conditions. These compounds were added to modified *Saccharomyces cerevisiae* synthetic complete (SC) medium (*SI Appendix*, Table S1) with different dissolution methods: fatty acid salts in aqueous solution, fatty acids and β-MAGs dissolved in ethanol, or fatty acids conjugated with bovine serum albumin (BSA). Fatty acid–BSA conjugates (C12:0, C14:0, C16:0, C18:0, and C18:1) were dissolved to a final concentration of 0.5 mM, whereas potassium salts of fatty acids (C12:0, C14:0, and C16:0) were converted into an insoluble form due to the formation of metal soap with Ca^2+^ and Mg^2+^ in SC medium. Moreover, fatty acids (C12:0, C14:0, C16:0, anteiso-C15:0, C16:1Δ9Z, and C16:1Δ11Z) and β-MAGs (C14:0 and C16:0) were aggregated in solid culture (*SI Appendix*, Fig. S1). Symbiotically generated spores of *R. irregularis* were used as starting material (parent spores) for asymbiotic culture. In the absence of fatty acids, no increase in biomass was detected for *R. irregularis* (Fig. *1A* and *B*). After spore germination, each *R. irregularis* spore generated a thick and short subtending hypha that soon branched to produce straight-growing thick hyphae (hereafter referred to as runner hyphae), with several thin lateral hyphae displaying a low level of ramification (Fig. 1*D* and *E*). Hyphal elongation ceased within one or two weeks after germination. By contrast, myristates (C14:0), unlike other tested chemicals, substantially increased fungal biomass regardless of the method used for their incorporation in the medium (Fig. 1*A*). Moreover, the increase in fungal biomass in the culture system along time depended on the amount of myristate (Fig. 1*B*). As a consequence, the total dry weight of a two-month-old colony treated with 1 mM potassium myristate was double that of parent spores *(SI Appendix*, Fig. S1*A*). Surprisingly, 16:0 β-MAG and palmitate (16:0), candidate compounds released from arbusculated host cells during symbiosis (18, 19), did not increase biomass production. However, during cultivation with myristate, vigorous hyphal development and subsequent differentiation of secondary spores were observed (Fig. *1F–P* and *SI Appendix*, Fig. S1). After germination, *R. irregularis* differentiated a few densely packed coil (DPC)-like structures from the runner hyphae in the vicinity of the parent spore (Fig. 1*G*). The DPC is an extensively branched hyphal structure, which was first observed in *R. irregularis* co-cultivated with *P. validus* (24) and whose existence was confirmed in fungal materials supplemented with palmitoleic acid and anteiso-C15:0 (11). Furthermore, *R. irregularis* elongated its runner hyphae by generating short-branched hyphae similar to branched absorbing structures (BAS) (25) and expanded its habitat by generating new runner hyphae (Fig. 1*H–J*). The branched hyphal structures (hereafter referred to as BAS), small bunches of short and thin branches, were generated from the runner hyphae at short intervals (Fig. 1*K*). Interestingly, at the beginning of the cultivation with a potassium myristate supplement, numerous precipitates of putative myristate salt were observed throughout the growth medium; however, the precipitates around actively growing hyphae were progressively solubilized (Fig. 1*H, I*, and *L*). Moreover, myristate-induced secondary spores were frequently observed along the runner hyphae in the vicinity of parent spores (Fig. *1M*). Myristate-induced spores also occurred apically or intercalary along the lateral branches of the extensively growing runner hyphae (Fig. 1*N* and *O*). These spores were approximately 50 μm in diameter, which is almost half the size of parent spores (Fig. *1F, M*, and *P*). In the presence of palmitoleic acid (C16:1Δ9Z) in the medium, extensive hyphal branching and secondary spore formation were observed, which is consistent with the results by Kameoka and co-workers (11). However, in our system hyphal growth was not associated with an increase in biomass by the application of palmitoleic acid (Fig. *1A* and *SI Appendix*, Fig. S1*E*). When lauric acid (C12:0) was applied as a lauric acid–BSA conjugate, *R. irregularis* showed active elongation of runner hyphae with few DPC, BAS, or secondary spores (*SI Appendix*, Fig. S1*F*). However, this conjugate did not significantly increase fungal biomass (Fig. 1*A*). Thus, as the lauric acid–BSA conjugate was effective on hyphal elongation even at low concentrations (1 and 10 μM), lauric acid is not likely to be utilized as a macronutrient for *R. irregularis (SI Appendix*, Fig. S1*H*).

**Fig. 1.**
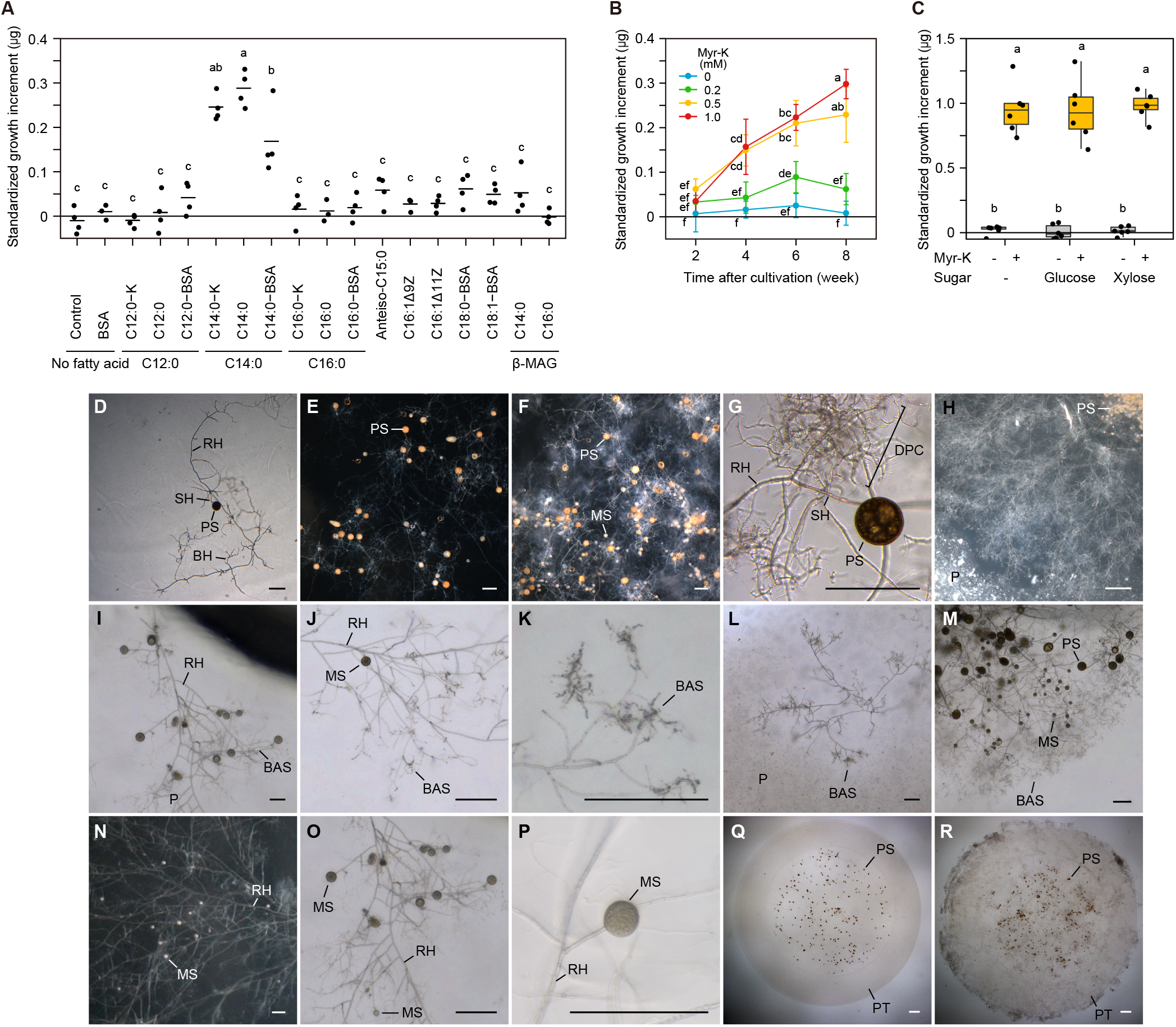
Asymbiotic culture of *R. irregularis* in the presence of fatty acids. (*A*) Standardized growth increment (see Materials and Methods) during eight weeks in the modified SC solid medium supplemented with potassium salts of fatty acids, fatty acids and β-MAGs, or fatty acid-BSA conjugates. Horizontal lines indicate mean values (*n* = 3–4). (*B*) Time course of biomass production at different amounts of potassium myristate (Myr-K). Error bars represent 95% confidence intervals (*n* = 5–6). (*C*) Biomass production in an immobilized cell culture system. Immobilized fungal spores were incubated in half-strength modified SC medium supplemented with combinations of potassium myristate and sugars after eight weeks of cultivation. The boxes show the first quartile, the median, and the third quartile; the whiskers reach to the 1.5× interquartile range, and data points for each treatment are displayed (*n* = 6). The same lowercase letter indicates no significant difference (Tukey’s test, *P* < 0.05; *A–C*). Fungal growth in the solid medium without fatty acids (*D* and *E*) or with potassium myristate (*F–P*). (*D*) Germinating spore. Hyphal elongation without fatty acids (*E*) and with potassium myristate (*F*) after eight weeks of cultivation. (*G*) DPC-like structures formed around a parent spore. (*H*) Radial growth of fungal mycelium. (*I*) Elongation of runner hyphae. (*J*) Branching of runner hyphae and formation of BAS. (*K*) Magnified image of BAS. (*L*) Front line of elongated mycelium. The medium around the fungal hyphae became transparent, indicating that the precipitates of metal soaps were solubilized. (*M*) Myristate-induced secondary spores generated around the parent spores. (*N* and *O*) Myristate-induced spores formed along the runner hyphae. (*P*) Magnified image of a myristate-induced spore. Fungal growth in the immobilized cell culture system without fatty acids (*Q*) and with potassium myristate (*R*) after eight weeks of cultivation. See *SI Appendix*, Table S2 for sample details. BAS, branched absorbing structure; BH, branching hypha; DPC, densely packed coil; MS, myristate-induced spore; P, precipitate of metal soaps; PS, parent spore; PT, Phytagel tablet; RH, runner hypha; and SH, subtending hypha. Scale bars: 200 μm (*D–G* and *I–P*) and 1,000 μm (*H, Q*, and *R*).

We further developed our culture system to promote fungal growth under asymbiotic conditions. First, we examined fungal growth in liquid culture with a modified SC medium containing potassium myristate. Myristate was effective in enhancing fungal biomass also in liquid culture (*SI Appendix*, Fig. S2*A*). After germination, precipitates of metal soaps attached to the hyphal surface (*SI Appendix*, Fig. S2*B*). Afterwards, as the fungal hyphae continued to elongate and became tangled, the metal soaps on the surface gradually disappeared. However, typical BAS were not recognized due to the aggregation of fungal hyphae, although highly branched fine hyphae were observed in the presence of myristate (*SI Appendix*, Fig. S2*C*). In contrast to solid culture, very few secondary spores formed in liquid culture. Next, an immobilized cell culture system was tested, in which an inoculum of *R. irregularis* spores embedded in the center of a Phytagel tablet was incubated in modified SC liquid medium (*SI Appendix*, Fig. S3*A*). This culture system can prevent the aggregation of fungal hyphae and facilitate the diffusion of medium components in the Phytagel tablets. Notably, AM fungal growth in the immobilized cell culture system with potassium myristate was increased by 3-4-fold with respect to that in solid and liquid cultures (Fig. *1A* and *C*; *SI Appendix*, Fig. S2*A*). Hyphal elongation was very active, and some hyphae even spread out of the Phytagel tablets and continued to grow (Fig. *1Q* and *R* and *SI Appendix*, Fig. S3*B–D*). The elongation pattern of hyphae was similar to that in solid culture. For instance, DPC-like structures, runner hyphae, BAS, and myristate-induced spores were observed (*SI Appendix*, Fig. S3*E–I*). Also, myristate-induced spores were still smaller than parent spores (*SI Appendix*, Fig. S3*J*).

As monosaccharides are utilized by AM fungi as carbon sources during symbiosis (12-14), we examined whether monosaccharides influenced on the growth of *R. irregularis* in an immobilized cell culture system. However, in the absence of potassium myristate, neither glucose nor xylose promoted biomass production (Fig. 1*C*). Similarly, no additional biomass increase was detected upon treatment with a combination of monosaccharides and potassium myristate. Modified SC medium contained 1 mM glycerol, which is a potential carbon source because *R. irregularis* can absorb and metabolize it under asymbiotic conditions (26). However, glycerol did not increase biomass production of *R. irregularis* with or without myristate (*SI Appendix*, Fig. S1*B–D*).

To ascertain whether myristate exerts similar effects in other AM fungal species, *Rhizophagus clarus* (Order: Glomerales, Family: Glomeraceae) and *Gigaspora margarita* (Order: Diversisporales, Family: Gigasporaceae), belonging to the same genus as and a different order than *R. irregularis*, respectively, were cultured in the same conditions. When incubated in a medium without fatty acids, *R. clarus* produced short runner hyphae from the parent spores and formed a few small secondary spores, although hyphal elongation soon ceased (*SI Appendix*, Fig. S4). Conversely, in the presence of myristate, *R. clarus* showed active hyphal growth and sporulation accompanied by an increase in biomass. After germination, runner hyphae were vigorously elongated and sometimes branched dichotomously. In addition, BAS-like structures were generated at regular intervals from several long runner hyphae. Myristate-induced secondary spores were also formed on short lateral branches deriving from runner hyphae. Similar to those seen in *R. irregularis*, the myristate-induced spores of *R. clarus* were half the size of the parent spores and possessed thinner spore walls. Since we could not obtain an adequate number of sterile spores of *G. margarita*, we only analyzed its growth pattern. *G. margarita* produced longer hyphae than *R. irregularis* and *R. clarus* even in the absence of myristate, possibly owing to its large spores with more abundant energy reserves (*SI Appendix*, Fig. S5). After transfer to a myristate-containing medium, *G. margarita* produced much longer runner hyphae, from which BAS-like structures were generated. No newly formed spores were observed in *G. margarita*, although auxiliary cells (subglobose, hyaline cells clustered in groups of approximately 10 with spiny ornamentations borne on coiled hyphae) differentiated in both media.

### Myristate-induced spores have infection capability

Myristate-induced spores of *R. irregularis* began to differentiate from two weeks after cultivation in solid medium. The number of myristate-induced spores increased with time and amount of myristate and finally reached 1.4 spores per parent spore eight weeks after treatment (Fig. 2*A*). However, these spores were still half the size of the parent spores (Fig. 2*B*). In the immobilized cell culture system, the number of *R. irregularis* myristate-induced spores was equal to that in solid culture, and addition of xylose further promoted their production, reaching up to 2.5 spores per parent spore (Fig. 2*C* and *SI Appendix*, Fig. S6). Myristate-induced spores were initially white to pale yellow and gradually turned to a yellow-brown color, similar to symbiotically generated spores (*SI Appendix*, Fig. S6*A* and Fig. S7*A*). Moreover, both spores induced by myristate and those produced symbiotically showed three spore wall layers originating from cylindrical subtending hyphae, although wall layers of myristate-induced spores were less thick than those of symbiotically generated spores (*SI Appendix*, Fig. S7*B*). Many nuclei, vacuoles, and lipid droplets were observed in both spore types (*SI Appendix*, Fig. S7*A* and *B*). To test whether myristate-induced spores could colonize plant roots, single spores generated in the immobilized cell culture system with potassium myristate and xylose were inoculated to carrot hairy roots. In total, 245 spores were examined in six independent experiments (*SI Appendix*, Table S3). Myristate-induced spores displayed infectivity towards hairy roots, triggering the production of next-generation mature spores on the extraradical hyphae that emerged from the roots (Fig. 2*D–G*). Approximately half of the germinated spores could colonize hairy roots and produce daughter spores, albeit large variations in the germination rate and infectivity of spores were observed among the trials due to the effect of experimental manipulations (*SI Appendix*, Table S3).

**Fig. 2.**
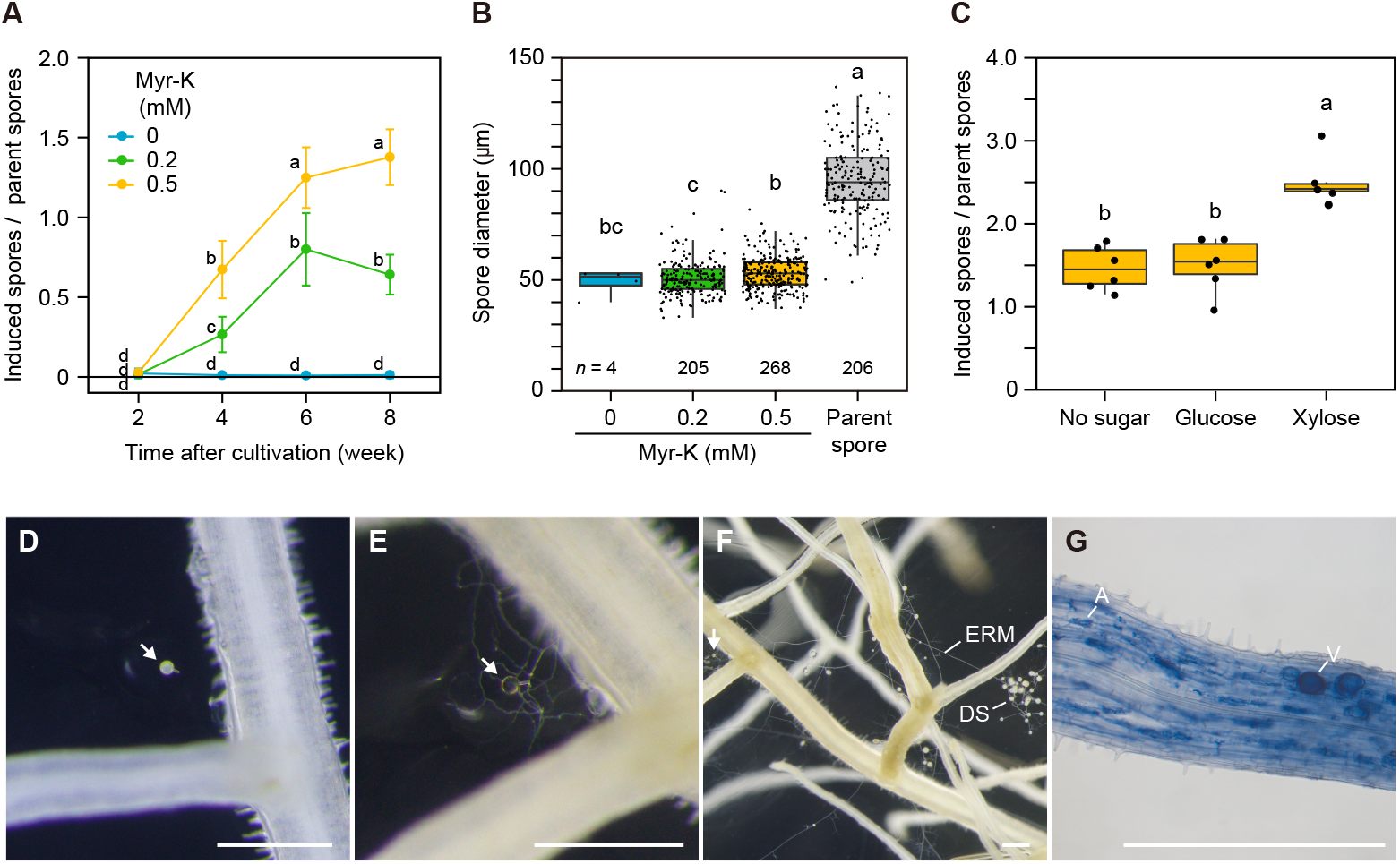
Spore formation of *R. irregularis* under asymbiotic conditions. Number (*A*) and diameter (*B*) of myristate-induced spores generated in solid medium containing different amounts of potassium myristate (Myr-K). Spore diameter was measured after eight weeks of cultivation. (*C*) Number of myristate-induced spores in an immobilized cell culture containing 0.5 mM potassium myristate and 5 mM monosaccharides after eight weeks of cultivation. Error bars in (*A*) represent 95% confidence intervals. For each boxplot, the boxes show the first quartile, the median, and the third quartile; the whiskers reach to the 1.5× interquartile range, and data points for each treatment are displayed. The same lowercase letter indicates no significant difference (Tukey’s test, *P* < 0.05, data were transformed as log_10_ (x+0.5), *n* = 3–6 (*A*); Steel-Dwass test, *P* < 0.05 (*B*); and Tukey’s test, *P* < 0.05, *n* = 6 (*C*)). (*D–G*) Inoculation of a single myristate-induced spore to carrot hairy roots. (*D*) Inoculated spore that was produced in the immobilized cell culture containing potassium myristate and xylose. (*E*) Germination of the inoculated spore. (*F*) Daughter spores (DS) on extraradical mycelia (ERM) emerged from the carrot hairy roots. (*G*) Colonization of the carrot hairy roots by *R. irregularis*. Arrows indicate an inoculated myristate-induced spore. A, arbuscule; and V, vesicle. See *SI Appendix*, Table S2 for sample details. Scale bars: 500 μm.

### AM fungi utilize myristate as a carbon and energy source

To address the use of fatty acids by AM fungi under asymbiotic conditions, we analyzed fatty acid uptake using two fluorescent fatty acid derivatives of different chain lengths, C1-BODIPY 500/510 C12 and BODIPY FL C16. *R. irregularis* absorbed C1-BODIPY 500/510 C_12_ through its BAS (not runner hyphae) within 10 min from first exposure (Fig. 3*A*). The probe was first taken up from the hyphal tips of BAS (Fig. 3*A*), then fluorescent signals were observed over time in lipid body-like structures (27, 28) within runner hyphae, which were translocated by cytoplasmic streaming (*SI Appendix*, Fig. S8*B* and Movie S1). BODIPY FL C_16_ was also absorbed by the BAS within 4 h, while faint probe signals were observed during a short period of 10 min after the exposure (*SI Appendix*, Fig. S9*A–C*). Long exposure to these probes resulted in signals localized in myristate-induced spores as well as BAS, runner hyphae, and DPC, but parent spores were rarely labeled (*SI Appendix*, Fig. S8*A–E* and Fig. S9*D–H*). Moreover, germ tubes deriving from germinating spores incubated in sterilized water also took up the fluorescently labeled fatty acids, indicating that the activation of AM fungal hyphae by myristate is not necessarily required for fatty acid uptake (*SI Appendix*, Fig. S8*F–G* and S9*I–J*). Gene expression analysis confirmed that potassium myristate transcriptionally activated β-oxidation, the glyoxylate cycle, gluconeogenesis, and the TCA cycle 3 h after its application (Fig. 3*B*). Interestingly, a gene encoding a *N*-myristoyl transferase (NMT), which catalyzes the myristoylation of proteins, was upregulated by myristate. To further assess whether myristate provides the carbon skeleton for fungal cell components, we applied [1-^13^C]myristate to the culture medium and analyzed the cell wall components of *R. irregularis*. After the extraction of fungal cell walls, chitin and chitosan were converted into glucosamine through acid hydrolysis. Liquid chromatography-mass spectrometry (LC-MS) analysis of the extracted glucosamine demonstrated that the relative ion intensity of M+1 was significantly higher in AM fungi supplemented with [1-^13^C]myristate (mean 16.4%) than in those supplemented with non-labeled myristate (6.1%) (Fig. 3*C*). This finding indicated that external myristate was taken up by *R. irregularis* and utilized for the biosynthesis of chitin and chitosan in fungal cell walls. In addition, to evaluate the use of myristate as an energy source, ATP production was measured after the application of myristate. In particular, we applied myristate to germinating hyphae and measured their ATP content 12 h after application. Notably, ATP content increased by 2.4-fold in the presence of myristate (Fig. 3*D*). In addition, when carbonyl cyanide *m*-chlorophenylhydrazone (CCCP), an inhibitor of mitochondrial membrane depolarization, was simultaneously applied to the hyphae, ATP content did not increase even after fatty acid application. The translocation of the fluorescent fatty acid probes to myristate-induced spores (*SI Appendix*, Fig. S8 and S9) prompted us to examine whether myristate is utilized to synthesize the major storage lipid, triacylglycerol (TAG). To this purpose, we cultured *R. irregularis* in medium supplemented with [1-^13^C]myristic acid. After extraction of lipids from the fungal materials, TAG was purified through preparative thin-layer chromatography (TLC) and analyzed by ^13^C-NMR. In the spectrum, two peaks at 173.1 and 173.4 ppm, which corresponded to the carboxyl carbons of acyl chains at the α and β positions, respectively, were observed at a much higher intensity than in the spectrum of TAG prepared from non-labeled fungal materials or monoxenically cultured *R. irregularis* (Fig. 3*E*). This finding showed that exogenous myristate was incorporated into TAG, especially in its acyl carboxyl components.

**Fig. 3.**
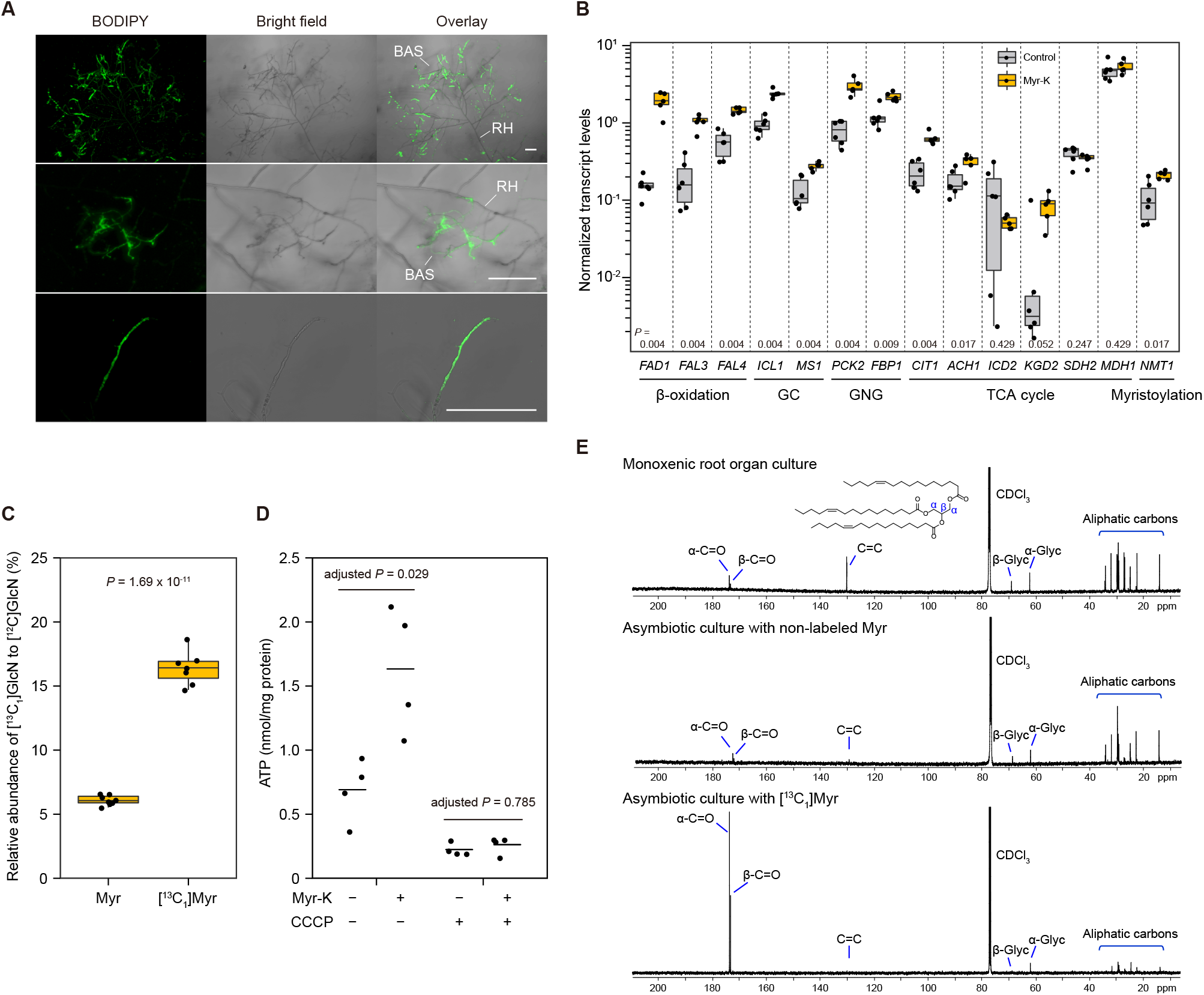
Utilization of fatty acids by *R. irregularis* under asymbiotic conditions. (*A*) Uptake of the fluorescently labeled fatty acid derivative C_1_-BODIPY 500/510 C_12_. A cluster of BAS (upper panels), magnified BAS (middle panels), and a single BAS hypha (lower panels). Optical sections captured using a confocal laser scanning microscope are projected. Fluorescence images and superimposed bright field images are shown. BAS, branched absorbing structure; and RH, runner hypha. Scale bars, 200 μm. (*B*) Gene expression analysis in *R. irregularis* 3 h after the application of potassium myristate (Myr-K) or water (control) shown by real-time RT-PCR. The *R. irregularis* elongation factor 1 beta and actin genes were used for normalization. For each boxplot, the boxes show the first quartile, the median, and the third quartile; the whiskers reach to the 1.5× interquartile range, and data points for each treatment are displayed (Myr-K: *n* = 5; control: *n* = 6). *P* values were calculated using the Wilcoxon–Mann–Whitney test. See *SI Appendix*, Table S4 for gene details. GC, glyoxylate cycle; GNG, gluconeogenesis. (*C*) Incorporation of carbon derived from exogenous myristate into cell wall components of *R. irregularis*. AM fungi were grown in the immobilized cell culture system and supplemented with 0.5 mM neutralized myristic acid (Myr) or [1-^13^C_1_]myristic acid ([^13^C_1_]Myr) for eight weeks. Glucosamine (GlcN) was extracted from the cell walls of the fungal materials without parent spores. The relative abundance of [^13^C_1_]GlcN ([M+1+H]^+^, *m/z* 181.19) to [^12^C]GlcN ([M+H]^+^, *m/z* 180.19) was calculated by LC-MS analysis. The reported *P* value is based on Student’s *t*-test (Myr: *n* = 8; [^13^C_1_]Myr: *n* = 7). (*D*) ATP content in *R. irregularis* 12 h after the application of potassium myristate or distilled water (control) in the presence or absence of CCCP. Horizontal lines indicate the mean values (*n* = 4). *P* values are based on Student’s *t*-test with Bonferroni correction. (*E*) ^13^C-NMR spectra of triacylglycerol (TAG) isolated from mycelial extracts of *R. irregularis* grown in monoxenic root organ culture (upper panel), asymbiotic culture supplemented with 1 mM neutralized myristic acid (middle panel), or [1-^13^C_1_]myristic acid (lower panel). Inset: chemical structure of a TAG. Experiments were repeated twice independently with similar results.

Since no signals denoting the presence of unsaturated TAG fatty acids in myristate-fed AM fungi were detected by ^13^C-NMR (Fig. 3*E*), the fatty acid composition is likely different from that in symbiotically generated spores. Further, AM fungi are predicted to obtain C16:0 β-MAG or palmitic acid from the host under symbiotic conditions (18, 19). Then, we analyzed the growth and lipid composition of AM fungi in the presence of a combination of myristate and C16:0 β-MAG or palmitate. We found that palmitate enhanced hyphal growth and biomass production of *R. irregularis* when combined with 0.5 mM myristate, but C16:0 β-MAG exerted no significant effect (Fig. 4*A* and *SI Appendix*, Fig. S10). Secondary spore formation was also stimulated by the combination of myristate and palmitate (Fig. 4*B*). In the presence of palmitate or C16:0 β-MAG together with myristate, spores displayed similar morphology but a slightly larger spore size than those induced by myristate (Fig. 4*C*; *SI Appendix*, Fig. S7 and Fig. S10). In addition, lipid droplets extracted from spores incubated in myristate seemed to be in a solid state, whereas those induced by a mixture of myristate and palmitate were liquid, similar to symbiotically generated spores (*SI Appendix*, Fig. S7*A*). This observation may reflect a change in the lipid composition of TAG. To prove this hypothesis, we analyzed the fatty acid composition (the major acyl group C14:0, C16:0, and C16:1Δ11) of AM fungal TAG by gas chromatography-mass spectrometry (GC-MS). In myristate-fed AM fungi, C14:0 was the dominant acyl group, while C16:0 and C16:1Δ11 were found in traces (Fig. 4*D*). Furthermore, [1-^13^C_1_]myristate labeling experiments showed that exogenous myristate was directly incorporated into TAG as C14:0 acyl groups (Fig. 4*E*). A significant fraction of C16:0 and C16:1Δ11 also derived from [1-^13^C_1_]myristate, indicating that myristate taken up into the fungus is elongated to C16:0 and desaturated into C16:1Δ11. In contrast, TAG in fungal materials supplemented with both myristate and palmitate contained high amounts of C16:0 and C16:1Δ11 (Fig. 4*D*). The majority of these acyl groups derived from the exogenous [1-^13^C_1_]palmitate (Fig. 4*E*). C16:1Δ11, a signature fatty acid for most AM fungi except those of the genera *Gigaspora, Archaeospora*, and *Paraglomus* (29), is likely to be generated from palmitate by the desaturase DES2, also known as OLE1-like, that is constitutively expressed in AM fungal hyphae (23, 30, 31). We next analyzed fungal cell wall components after incubation in a mixture of myristate and palmitate. The relative ion intensity of [^13^C_1_]glucosamine from AM fungi supplemented with [1-^13^C_1_]palmitate and non-labeled myristate was similar to the ion intensity of a non-labeled fatty acid mixture, indicating that carbon from exogenous palmitate was not incorporated into chitin and chitosan of fungal cell walls (Fig. 4*F*). Conversely, myristate-derived carbon was used for cell wall biosynthesis when [1-^13^C_1_]myristate and non-labeled palmitate were supplied.

**Fig. 4.**
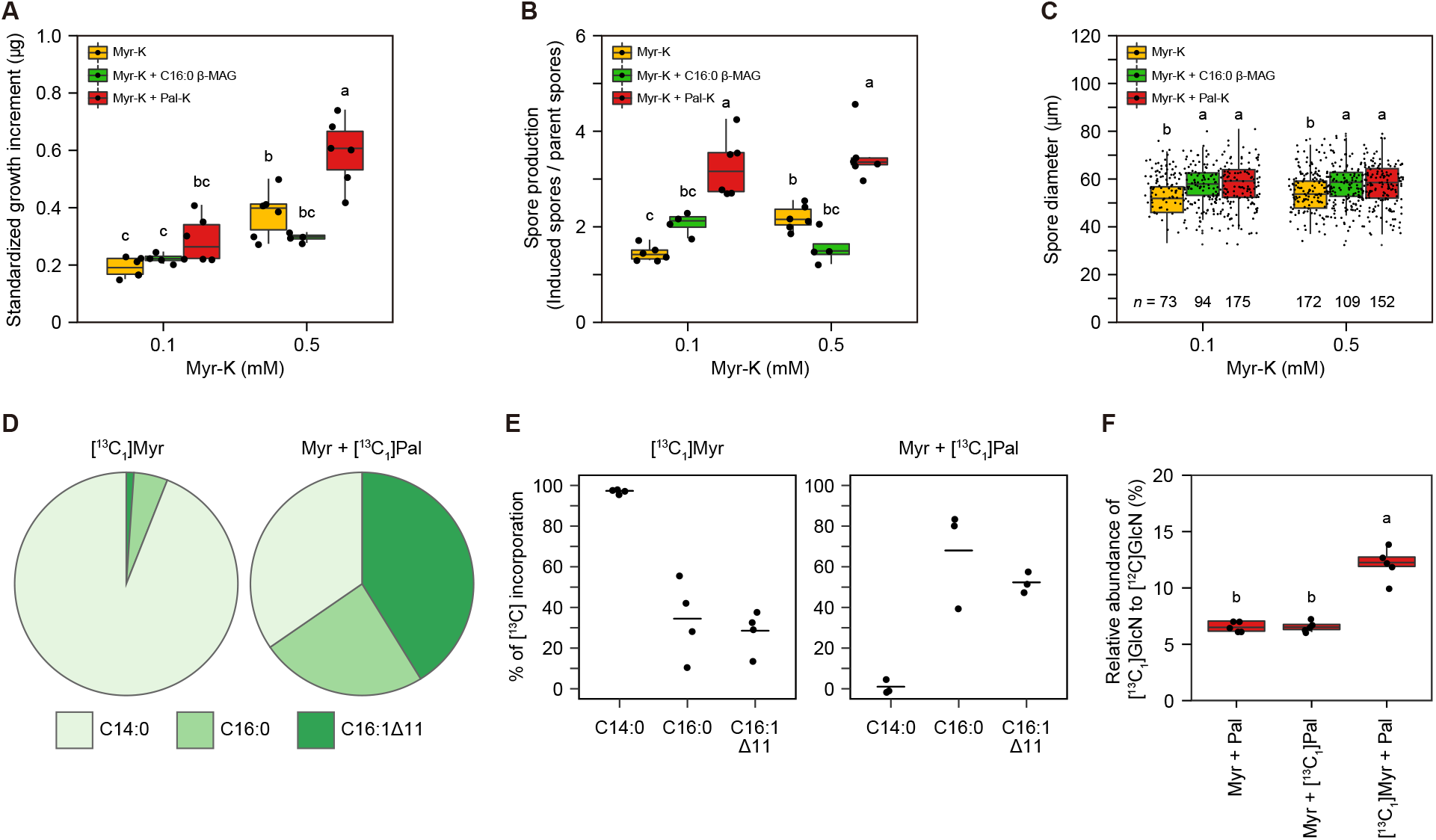
Effects of fatty acid mixtures on the growth and sporulation of *R. irregularis*. AM fungi were cultured in the immobilized cell culture system and supplemented with potassium myristate (0.1 or 0.5 mM Myr-K), either alone or in combination with 0.5 mM C16:0 *sn*-2 monoacylglycerols (β-MAG C16:0) or potassium palmitate (Pal-K) (*n* = 4–6). See *SI Appendix*, Table S2 for sample details. Standardized growth increment (*A*) and number (*B*) and diameter (*C*) of myristate-induced secondary spores after eight weeks of cultivation. For each boxplot, the boxes show the first quartile, the median, and the third quartile; the whiskers reach to the 1.5× interquartile range, and data points for each treatment are displayed. The same lowercase letter indicates that there is no significant difference (Tukey’s test, *P* < 0.05). (*D*) Composition of C14:0, C16:0, and C16:1Δ11 fatty acids of triacylglycerol (TAG) isolated from fungal materials grown in asymbiotic culture and supplemented with 0.5 mM neutralized [1-^13^C_1_]myristic acid ([^13^C_1_]Myr) or 0.1 mM potassium myristate plus 0.5 mM neutralized [1-^13^C_1_]palmitic acid (Myr + [^13^C_1_]Pal). (*E*) Percentage of [^13^C], derived from the labeled myristate or palmitate, incorporated into TAG. Horizontal lines indicate mean values (*n* = 3–4). (*F*) Incorporation of carbon derived from exogenous myristate and palmitate into cell wall components of *R. irregularis*. AM fungi were supplemented with labeled or non-labeled, neutralized myristic acid (0.5 mM) and palmitic acid (0.5 mM). Glucosamine (GlcN) was extracted from fungal cell walls without parent spores. Relative abundance of [^13^C_1_]GlcN ([M+1+H]^+^, *m/z* 181.19) to [^12^C]GlcN ([M+H]^+^, *m/z* 180.19) was calculated using data from LC-MS analysis. The same lowercase letter indicates no significant difference (Tukey’s test, *P* < 0.05, *n* = 5).

## Discussion

AM fungi have an obligate biotrophic lifestyle, i.e., these fungi depend on host-derived nutrients for their growth. Recently, AM fungi have been shown to receive lipids from the host via arbuscules (1-3, 18, 19); however, how these fungi utilize these lipids as nutrients is largely unknown. Here we show that myristate (C14:0) can be used as a carbon and energy source for the hyphal growth of *R. irregularis* under asymbiotic conditions, thereby providing the first evidence of increasing AM fungal biomass in a pure culture system. Myristate also promoted the growth of *R. clarus* and *G. margarita*, suggesting that myristate is effective in promoting asymbiotic growth in a wider range of AM fungal species. *R. irregularis* and *R. clarus* elongated their hyphae and formed secondary spores in a similar manner. In contrast, no secondary spores were observed in *G. margarita*, although long runner hyphae with BAS were generated in the presence of myristate. Considerable variation in the fatty acid composition of spores was observed between the genera *Rhizophagus* and *Gigaspora* (20, 29); therefore, their utilization of and response to exogenous fatty acids might also differ. Notably, we confirmed that myristate-induced spores of *R. irregularis* are infective propagules capable of generating symbiotic daughter spores, as previously demonstrated for palmitoleic acid-induced secondary spores (11). Moreover, the observation that fluorescently labeled fatty acid derivatives of different chain length accumulated in *R. irregularis* hyphae suggested that AM fungi are likely to absorb fatty acids non-specifically. However, myristate was the only fatty acid effective in promoting biomass production under our culture conditions. This is a surprising finding because 16:0 β-MAG or its related chemicals are thought to be transferred from the host to AM fungi under symbiotic conditions (18, 19). Meanwhile, as the plant enzymes FatM and RAM2, responsible for AM-specific lipid biosynthesis, have been found to use C14:0-containing molecular species as well as C16:0 as a substrate *in vitro* (3, 19), and myristate and C14:0 α-MAG have been detected in mycorrhizal roots and fungal spores, albeit in very small amounts (18, 20, 29), it is plausible that myristate is provided to AM fungi by the host. Myristate may be essential for the biological processes of AM fungi via myristate-specific metabolic pathways. Indeed, myristate is used for the lipid modification of proteins, called protein *N*-myristoylation, in a variety of eukaryotes (32-34). Protein *N*-myristoylation is catalyzed by NMT, which transfers myristate from myristoyl-CoA to the N-terminal glycine residue of target proteins (35). *N*-myristoylated proteins are involved in diverse cellular processes such as protein phosphorylation and signal transduction. Interestingly, disruption of the *NMT* gene causes recessive lethality in several fungal species (36-38). As indicated by the upregulation of the *R. irregularis NMT1* gene upon application of myristate, myristate may participate in boosting AM fungal growth by inducing protein *N*-myristoylation as well as by acting as a carbon source. If this was the case, we predict that fatty acids other than myristate would be also used for fungal growth if enough myristate was provided to AM fungi to induce *N*-myristoylation. Consistently, a mixture of myristate and palmitate further promoted fungal growth and led to a marked increase in the proportion of C16:0 and C16:1Δ11 acyl groups in fungal TAG, which are the predominant ones in symbiotically grown *R. irregularis* (4, 39). However, carbon derived from exogenous palmitate was not used for cell wall biosynthesis. On the other hand, in the presence of myristate, palmitate might stimulate the metabolism of stored lipids in parent spores for asymbiotic growth or serve as an energy source. The former hypothesis is consistent with the results of our [1-^13^C_1_]palmitate labeling experiment, in which exogenous [1-^13^C_1_]palmitate and non-labeled palmitate, likely derived from stored lipids, were incorporated into lipids of newly produced hyphae and secondary spores. However, palmitate might become available as a carbon source for AM fungal growth under both symbiotic and asymbiotic conditions in the presence of factors that were not considered in our culture system. Future studies may better elucidate how AM fungi use different types of fatty acids.

Fungal cell proliferation requires a sufficient supply of nutrients. Particularly, carbon sources are critical for producing the building blocks of cells and generating ATP. The observed incorporation of ^13^C in TAG and cell wall components of *R. irregularis* when [1-^13^C]myristate was supplied indicated that myristate is utilized as a carbon source to synthesize cellular components. During symbiosis, β-oxidation, the glyoxylate cycle, and gluconeogenesis are active in AM fungi, and these metabolic pathways have been proposed to play a crucial role in the generation of carbohydrates from lipids (14, 26, 40, 41). Since the expression of major genes involved in β-oxidation, the glyoxylate cycle, gluconeogenesis, and the TCA cycle was induced by the application of myristate, absorbed myristate is likely metabolized through these metabolic pathways and the resulting carbohydrates are used to build the backbone of fungal cells. Furthermore, increased ATP levels upon myristate addition indicated that myristate can also be used as an electron donor for the respiration of AM fungi. In our culture systems, hexoses did not induce an increase in fungal biomass, although xylose in combination with myristate triggered the production of a great number of secondary spores. This finding implies that AM fungi can hardly use sugars as carbon sources under these conditions. This observation is consistent with the results of previous ^13^C labeling experiments, which demonstrated a reduced hexose uptake by germinating spores than by intraradical hyphae (40). However, hexoses derived from the host are taken up via fungal monosaccharide transporters in intraradical hyphae and/or arbuscules during symbiosis (13). In addition, the monosaccharide transporter *MST2* gene is expressed even in BAS formed on the medium in a carrot hairy root system (23) and induced in extraradical hyphae treated with xylose (16). Thus, we cannot rule out the possibility that AM fungi can utilize exogenous sugars for their growth under asymbiotic conditions.

A characteristic response of AM fungi to myristate is the branching of runner hyphae and the formation of BAS. To date, a number of chemical compounds, such as strigolactones, 2-hydroxy fatty acids, palmitoleic acid, and branched fatty acids, have been found to induce hyphal branching during the presymbiotic phase (11, 42, 43). For example, strigolactones stimulate 5–6^th^-order hyphal branching in *G. margarita* at extremely low concentrations (42), but they only moderately induce hyphal branching in *Rhizophagus* sp. LPA8 (44). Moreover, two 2-hydroxy fatty acids, 2OH-C14:0 and 2OH-C12:0, also affected presymbiotic hyphal growth in *Gigaspora* spp., and these AM fungi were found to produce multiple lateral branches along the primary germ tubes; however, 2OH-C14:0 and 2OH-C12:0 did not elicit any morphological change in *R. irregularis* (43). In contrast, palmitoleic acid and branched fatty acids stimulate hyphal branching of *R. irregularis* and *R. clarus* at low concentrations, but they do not display this effect in *G. margarita* (11). In particular, palmitoleic acid was found to induce high-degree short branching and sporulation. However, the effect of myristate on hyphal branching and elongation was completely different from the effects of these known stimulants. Indeed, we observed that myristate induced extensive hyphal branching in *R. irregularis, R. clarus*, and *G. margarita* at high concentrations. In contrast, it is known that these AM fungi do not respond to low concentrations of myristate (11, 43). To explain this phenomenon, it is likely that myristate is utilized as a nutrient, with subsequent activation of fungal metabolism and gene expression, which results in the differentiation of BAS, DPC, and branched runner hyphae to further absorb myristate. Indeed, we observed that fatty acids were taken up by the fungal BAS and DPC. Then, absorbed fatty acids or their metabolites are likely to be stored in lipid bodies and translocated to runner hyphae, as previously observed in symbiotic extraradical hyphae (26, 28, 41). Part of the fatty acids or lipids might be also delivered to newly formed spores, where they may be used for spore germination and subsequent hyphal elongation.

In conclusion, asymbiotic growth of AM fungi can be supported by externally supplied fatty acids, leadings to the possibility of generating a pure culture of biotrophic AM fungi. Although myristate initiates AM fungal growth and sporulation, the size of myristate-induced spores remains small compared with that of symbiotically generated spores. Similar results were obtained for palmitoleic acid-induced spores (11). As smaller spores show low germination rates and infectivity (45), spore maturation in the absence of the host represents an exciting future challenge for the pure culture of AM fungi. Altogether, our findings have shed new light on the cellular and molecular biology of AM fungi and carry important implications for the development of new strategies for the genetic transformation and the production of inocula of these organisms.

## Materials and Methods

Materials and methods used in this study are described in detail in *SI Appendix, Extended Methods*.

### Biological materials

Sterile spore suspensions of the AM fungus *R. irregularis* DAOM197198 (or DAOM181602, another voucher number for the same fungus) were purchased from Premier Tech. Hyphae included in the spore suspension were removed by density-gradient centrifugation using gastrografin as described in *SI Appendix, Extended Methods. R. clarus* HR1 (MAFF520076) and *G. margarita* K-1 (MAFF520052) were also used for asymbiotic culture.

### Asymbiotic culture

Approximately 300–400 parent spores of *R. irregularis* were placed on 0.3% Phytagel (Sigma-Aldrich) plates containing modified SC medium (*SI Appendix*, Table S1) and then covered with 0.3% Phytagel dissolved in 3 mM magnesium sulfate in a 12-well culture plate. For liquid culture, Phytagel was removed from the medium. Three types of fatty acids were added to the medium: fatty acid salts, fatty acids in an organic solvent, or fatty acids conjugated with BSA. The plates were incubated at 28 °C in the dark. Hyphal elongation was observed under a dissecting microscope and light microscopes.

### Immobilized cell culture

An overview of the immobilized cell culture system is represented in the *SI Appendix*, Fig. S3*A*. Six-mm high and 17.5-mm wide Phytagel tablets, with 3-mm deep and 11.5-mm wide circular incisions containing *R. irregularis* spores, were transferred into a 6-well culture plate. Each well was filled with 5 mL of full- or half-strength modified SC liquid medium with an appropriate amount of fatty acids and monosaccharides. AM fungi were grown at 28 °C in the dark. During the culture period, the liquid medium was changed once a month.

### Asymbiotic culture of *R. clarus* and *G. margarita*

Sterile spores of *R. clarus* were prepared using a monoxenic system with carrot hairy roots. *G. margarita* spores were extracted from soil in pot culture and sterilized using chloramine T. These AM fungi were cultured in Phytagel covered with half-strength modified SC medium containing 0.5 mM potassium myristate at 28 °C for 12 weeks.

### Measurement of fungal biomass

Fungal materials were recovered from gels in wells of a culture plate by melting the gels in citrate buffer and weighed with a micro analytical balance. The number of parent spores in each well was counted in advance under a dissecting microscope. The standardized growth increment of AM fungi was calculated by dividing the total fungal dry weight in each well by the number of parent spores and subtracting the mean dry weight of a parent spore.

### Spore morphology

Spores were mounted with polyvinyl alcohol–lactic acid–glycerol (PVLG) or Melzer’s reagent for microscopic observation. Spores were incubated with 10 μM SYTO 13 Green Fluorescent Nucleic Acid Stain (Thermo Fisher Scientific) for 2 h and observed by epifluorescence microscopy. Transmission electron microscopy was performed to analyze the ultrastructure of spores according to Kameoka and co-workers (11).

### Single spore inoculation

A single myristate-induced spore produced in the immobilized cell culture system in a half-strength modified SC medium supplemented with 0.5 mM potassium myristate and 5 mM xylose was placed onto plates with carrot hairy roots using a pipette. The production of daughter spores on extraradical hyphae emerging from hairy roots was observed under a dissecting microscope. AM fungal colonization was confirmed by trypan blue staining.

### Fatty acid uptake

*R. irregularis* was grown in an immobilized cell culture system with modified SC medium containing 0.5 mM potassium myristate for six to eight weeks. Fungal hyphae were stained with 0.5 mM 4.4-difluoro-5-methyl-4-bora-3a,4a-diaza-*s*-indacene-3-dodecanoic acid (C_1_-BODIPY 500/510 C_12_, Thermo Fisher Scientific) or 4.4-difluoro-5,7-dimethyl-4-bora-3a,4a-diaza-*s*-indacene-3-hexadecanoic acid (BODIPY FL C_16_, Thermo Fisher Scientific) in a modified SC medium. After a 10-minute or 4-h incubation, fungal hyphae protruding outside a Phytagel tablet were observed using a laser scanning confocal microscope or an epifluorescence microscope. For the samples incubated for over one day in the medium containing the fluorescent probes, a Phytagel tablet containing fungal materials was melted by adding citrate buffer. Fluorescent signals were observed under an epifluorescence microscope. We also assayed fatty acid uptake by germ tubes grown in the absence of myristate.

### LC-MS analysis of glucosamine

*R. irregularis* was incubated for eight weeks in half-strength modified SC medium with one of the following five supplements: 1) 0.5 mM non-labeled myristic acid, 2) 0.5 mM [1-^13^C_1_]myristic acid (Taiyo Nippon Sanso), 3) 0.5 mM myristic acid and 0.5 mM palmitic acid, 4) 0.5 mM myristic acid and 0.5 mM [1-^13^C_1_]palmitic acid (Taiyo Nippon Sanso), and 5) 0.5 mM [1-^13^C_1_]myristic acid and 0.5 mM palmitic acid. All fatty acids used were neutralized in 200 mM potassium hydroxide. Extraction of glucosamine from fungal biomass and LC-MS analysis are described in *SI Appendix, Extended Methods*. The relative intensities of the molecular ion peaks of glucosamine ([M+H]^+^, *m/z* 180.19; and [M+1+H]^+^, *m/z* 181.19) were monitored. The relative fraction of M+1 with respect to that of M+0 in the glucosamine standard solution was 6.8%.

### ^13^C-NMR analysis of TAG

*R. irregularis* was cultured in modified SC solid medium supplied with 1 mM neutralized [1-^13^C]myristic acid (Cambridge Isotope Laboratories, Inc.) for 2.5 months. Extraction of lipids from fungal biomass, purification of TAG, and ^13^C-NMR analysis are described in *SI Appendix, Extended Methods*.

### GC-MS analysis of TAG

*R. irregularis* was incubated for eight weeks in half-strength modified SC medium with one of the following four supplements: 1) 0.5 mM neutralized myristic acid, 2) 0.5 mM neutralized [1-^13^C_1_]myristic acid, 3) 0.1 mM potassium myristate and 0.5 mM neutralized palmitic acid, and 4) 0.1 mM potassium myristate and 0.5 mM neutralized [1-^13^C_1_]palmitic acid. The extraction of lipids from fungal biomass, purification of TAG, and GC-MS analysis are described in *SI Appendix, Extended Methods*.

### Determination of ATP content

*R. irregularis* spores were incubated in sterilized water at 28 °C for five days. Potassium myristate was added to the germinating spores at a final concentration of 0.5 mM. For the control, the protonophore CCCP was simultaneously added at a final concentration of 50 μM. After incubation for 12 h, fungal materials were crushed in phosphate-buffered saline (PBS; pH 7.4) using a bead crusher (μT-12, TAITEC). ATP concentration was measured using the CellTiter-Glo Luminescent Cell Viability Assay kit (Promega). Protein concentration was assayed using the Qubit Protein Assay Kit (Thermo Fisher Scientific). ATP content in the germinating spores was calculated in nmol mg^-1^ of protein.

### Quantitative RT-PCR

*R. irregularis* was grown in an immobilized cell culture system with modified SC medium containing 0.5 mM potassium myristate for three weeks. Subsequently, AM fungi were incubated in the absence of fatty acids for 11 days to induce fatty acid starvation. After starvation, potassium myristate to a final concentration of 0.5 mM or sterilized water was added to the samples. After a 3-h incubation, fungal hyphae protruding outside a Phytagel tablet were recovered using forceps. RNA extraction, purification, cDNA synthesis, and semiquantitative PCR were conducted as described in *SI Appendix, Extended Methods*.

### Statistical analysis

All statistical analyses were performed using R version 3.5.2. Levene’s tests were applied to check for heteroscedasticity between treatment groups. Data were transformed as log_10_ (x + 0.5) where necessary. To examine the differences among experimental groups, data were analyzed with Student’s *t*-test, Tukey’s HSD test, Wilcoxon–Mann–Whitney test, and Steel–Dwass test, as appropriate. Differences at *P* < 0.05 were considered significant.

### Data Availability

All data used in the study are included in the paper and *SI Appendix*. All protocols are described in Materials and Methods and *SI Appendix, Extended Methods* or in cited references. If additional information is needed, it will be available upon request from the corresponding author.

## ACKNOWLEDGMENTS

We would like to thank T. Hagiwara, R. Oguchi, and R. Okita (Shinshu University) for their technical assistance and Enago and Editage for the English language review. LC-MS analysis and laser scanning confocal microscopy were conducted at the Research Center for Supports to Advanced Science, Shinshu University. This work was supported by ACCEL (JPMJAC1403) from the Japan Science and Technology Agency (to M.S., K.M., K.A, and K.S.) and the Grant-in-Aid for Scientific Research (15H01751 and 19H02861) from the Japan Society for the Promotion of Science (to K.S.).

