## Supplementary Information for "Myristate can be used as a carbon and energy source for the asymbiotic growth of arbuscular mycorrhizal fungi"

*SI Appendix*

**This PDF file includes:**

Extended Methods
Figs. S1 to S10
Tables S1 to S4
Captions for movie S1

**Other supplementary materials for this manuscript include the following:**

Movie S1

***Extended Methods***

**Biological materials.** Sterile spore suspensions of the arbuscular mycorrhizal (AM) fungus *Rhizophagus irregularis* DAOM197198 (or DAOM181602, another voucher number for the same fungus) were purchased from Premier Tech. Hyphae included in the spore suspension were removed by density-gradient centrifugation using gastrografin (972 mM amidotrizoate, 815 mM meglumine, and 157 mM sodium hydroxide). Five mL of 8%, 16%, 32%, and 48% gastrografin solution was carefully poured to form layers in a 50-mL centrifuge tube, and then 20 mL of spore suspension (approximately 4,000 spores $mL^{-1}$) was layered on top of the gastrografin solution. After centrifugation at 3,500 rpm for 15 min at room temperature using a swing rotor, middle layers containing spores were collected and transferred to a new tube. The collected layers were diluted with sterilized water and then centrifuged again. After the removal of the supernatant, spores were suspended in sterilized water, adjusted to approximately 10,000 spores $mL^{-1}$ and stored at 4 °C for later use. *Rhizophagus clarus* HR1 (or MAFF520076, another voucher number for the same fungus in the NARO Genebank; https://www.gene.affrc.go.jp/index_en.php) that was cultured monoxenically with *Agrobacterium rhizogenes*-induced hairy roots of carrot (1) and *Gigaspora margarita* K-1 (or MAFF520052 in the NARO Genebank) (2) were also used for asymbiotic culture.

**Asymbiotic culture.** Approximately 300–400 parent spores of *R. irregularis* were placed on 0.3% Phytagel (Sigma-Aldrich) plates containing modified SC medium (1.7 g $L^{-1}$ yeast nitrogen base [MP Biomedicals], 1.0 g $L^{-1}$ complete SC mixture [Formedium], 5 mM ammonium sulfate, 1 mM glycerol, 5 mg $L^{-1}$ thiamine hydrochloride, 5 mg $L^{-1}$ nicotinic acid, 5 mg $L^{-1}$ pyridoxal phosphate, and appropriate amounts of fatty acids) and then covered with 0.3% Phytagel dissolved in 3 mM magnesium sulfate in a 12-well culture plate. For liquid culture, Phytagel was removed from the medium. All fatty acids and fatty acid salts were obtained from Sigma-Aldrich (12-methyltetradecanoic acid [anteiso-C15:0]), MATREYA (11-hexadecenoic acid [C16:1Δ11Z]), Nacalai Tesque (palmitoleic acid [C16:1Δ9Z]), Tokyo Chemical Industry (lauric acid [C12:0], stearic acid [C18:0], and oleic acid [C18:1Δ9Z]), and Wako Pure Chemical Industries (myristic acid [C14:0], palmitic acid [C16:0], potassium laurate [C12:0-K], potassium myristate [C14:0-K], and potassium palmitate [C16:0-K]). Two β-monoacylglycerols, 2-myristoylglycerol and 2-palmitoylglycerol, were purchased from Santa Cruz Biotechnology. Three types of fatty acids were added to the medium: fatty acid salts, fatty acids in an organic solvent, or fatty acids conjugated with bovine serum albumin (BSA). Stock solutions of 100 mM free fatty acids and potassium salts of fatty acids were prepared by dissolving in ethanol and distilled water, respectively. For the preparation of fatty acid–BSA conjugates, an equal amount of 100 mM fatty acid was dissolved in ethanol, and 20% fatty acid-free BSA (Sigma-Aldrich) solution was mixed well and added with distilled water to generate 10 mM final concentration of fatty acid. More BSA solution was added to solubilize stearic acid. The plates were incubated at 28 °C in the dark. Hyphal elongation was observed under the dissecting microscope Stemi 508 (Carl Zeiss) and light microscope Primovert (Carl Zeiss). Digital images were captured with a digital CCD camera AxioCam MRc5 (Carl Zeiss) operated with AxioVision (Carl Zeiss). Fully focused images were captured on the microscope with BZ-X800 (Keyence).

**Immobilized cell culture.** An overview of the immobilized cell culture system is represented in the *SI Appendix*, Fig. S3*A*. Thirty-five mL of 0.75% Phytagel containing 3 mM magnesium sulfate was poured into a 90-mm Petri dish and solidified. Six-mm high and 17.5-mm wide Phytagel tablets, with 3-mm deep and 11.5-mm wide circular incisions, were cut out using a sterile double cork borer (inner diameter: 11.5 and 17.5 mm, respectively). The gel within the circular incision on the top side of the gel tablet was removed to 3 mm deep using a spatula or disposable pipette tip connected to an aspirator to prepare a hole. To flatten the bottom of the hole, a small amount of 0.75% Phytagel was added. Approximately 300–400 *R. irregularis* spores were placed in the hole and covered with 0.75% Phytagel containing 3 mM magnesium sulfate. The Phytagel tablets containing spores were transferred into a 6-well culture plate. Each well was filled with 5 mL of full- or half-strength modified SC liquid medium with an appropriate amount of fatty acids and monosaccharides. AM fungi were grown at 28 °C in the dark. During the culture period, the liquid medium was changed once a month.

**Asymbiotic culture of *R. clarus* and *G. margarita*.** Approximately 50 sterile spores of *R. clarus*, which were produced in a monoxenic system with carrot hairy roots, were placed in the center hole of 0.3% Phytagel, which was solidified in a 60-mm Petri dish. The hole was filled with 0.3% melting Phytagel with 3 mM magnesium sulfate. After solidification, the Phytagel was covered with 5 mL of a half-strength modified SC medium with 0.5 mM potassium myristate. Furthermore, *G. margarita* spores were extracted from soil in pot culture by wet-sieving, and they were sterilized using chloramine T following Cranenbrouck and co-workers (3). These spores were germinated on 0.75% Phytagel for 1 week. A single germinating spore was recovered with Phytagel using a sterile cork borer and transferred into a new 90-mm Petri dish. This dish was filled with 0.75% melting Phytagel with 3 mM magnesium sulfate. After solidification, the Phytagel was covered with 15 mL of half-strength modified SC medium supplemented with 0.5 mM potassium myristate. AM fungi were grown at 28 °C for 12 weeks. During the culture period, the liquid medium was not changed.

**Measurement of fungal biomass.** Gels containing fungal materials in wells of a culture plate were cut using a knife. A 1.5× volume of citrate buffer (8.3 mM trisodium citrate, 1.7 mM citric acid, and 1% Triton X-100; pH 6.0) was added to the well and mixed using a spatula. The plate was incubated at 50 °C for 20 min. The melted gel, including fungal materials, was transferred to a 5-mL tube and further incubated at 50 °C for 20 min. The sample was centrifuged at 3,500 rpm for 10 min at room temperature using a swing rotor. After the removal of the supernatant, approximately 5 mL of the citrate buffer was added, incubated at 50 °C for 10 min, and then centrifuged again. This step was repeated once. Fungal pellets were suspended in the remaining 1 mL of the citrate buffer and transferred to an antistatic 1.5-mL tube. After centrifugation at 3,500 rpm for 10 min, the supernatant was removed. The fungal pellet was washed with 1 mL of distilled water by centrifuging at 3,500 rpm for 10 min three times. After centrifugation at 12,000 rpm for 10 min using an angle rotor, the remaining water was completely removed. The pellet was dried at 70 °C for 48 h. After cooling down in a desiccator, fungal materials were weighed with a micro analytical balance (BM-252, A&D). The number of parent spores in each well was counted in advance under a dissecting microscope. The standardized growth increment of *R. irregularis* was calculated as follows:

$$\text{Standardized growth increment (µg)} = DW_{\text{t}}/N_{\text{p}} - DW_{\text{p}}$$

where $DW_t$ is the total fungal dry weight in each well after eight weeks of cultivation, $N_p$ is the number of parent spores, and $DW_p$ is the dry weight of a single parent spore. $DW_p$ was the mean value calculated from eight independent measurements in which total dry weight of several hundred spores was divided by the number of spores (*SI Appendix*, Fig. S1*A*).

**Spore morphology.** Spores were mounted with polyvinyl alcohol–lactic acid–glycerol (PVLG) or Melzer's reagent for microscopic observation using a light microscope (Axio Imager D1 microscope, Carl Zeiss). Spores were incubated with 10 µM SYTO 13 Green Fluorescent Nucleic Acid Stain (Thermo Fisher Scientific) for 2 h and observed by epifluorescence microscopy (Axio Imager D1 microscope). The fluorescence was excited with the band-pass filter BP470/40, and emitted fluorescence was detected with BP525/50. Transmission electron microscopy was performed to analyze the ultrastructure of spores according to Kameoka and co-workers (4).

**Single spore inoculation.** A single myristate-induced spore produced in the immobilized cell culture system in a half-strength modified SC medium supplemented with 0.5 mM potassium myristate and 5 mM xylose was placed onto plates with carrot hairy roots using a pipette according to Kameoka and co-workers (4). The production of daughter spores on extraradical hyphae emerging from hairy roots was observed under a dissecting microscope. AM fungal colonization was confirmed by trypan blue staining (5).

**Fatty acid uptake.** Fatty acid uptake was evaluated using fluorescent fatty acid analogs, 4,4-difluoro-5-methyl-4-bora-3a,4a-diaza-*s*-indacene-3-dodecanoic acid ($C_1$-BODIPY 500/510 $C_{12}$, Thermo Fisher Scientific) and 4,4-difluoro-5,7-dimethyl-4-bora-3a,4a-diaza-*s*-indacene-3-hexadecanoic acid (BODIPY FL $C_{16}$, Thermo Fisher Scientific). *R. irregularis* was grown in an immobilized cell culture system with modified SC medium containing 0.5 mM potassium myristate for six to eight weeks. After washing with the medium, AM fungi were incubated in modified SC medium with 0.5 mM fluorescent fatty acid analogs. After a 10-minute or 4-h incubation, fungal hyphae protruding outside a Phytagel tablet were observed using a laser scanning confocal microscope (Leica TCS SP2) or an epifluorescence microscope (Axio Imager D1 microscope). For the samples incubated for over one day in the medium containing the fluorescent probes, a Phytagel tablet containing fungal materials was melted by adding citrate buffer. Fluorescent signals were observed under the epifluorescence microscope. We also assayed fatty acid uptake by germ tubes grown in the absence of myristate. Spores of *R. irregularis* were germinated in sterile water for one week at 28 °C. Fluorescent probes were added at a final concentration of 0.5 mM. After incubating, fluorescent signals were observed using the epifluorescence microscope. For laser scanning confocal microscopy, $C_1$-BODIPY 500/510 $C_{12}$ was excited using a 488-nm wavelength argon laser, and fluorescence between 500 and 550 nm was detected. For epifluorescence microscopy, the fluorescent fatty acid analogs were excited with the band-pass filter BP470/40, and emitted fluorescence was detected with BP525/50. Digital images were captured with the digital CCD camera AxioCam MRm (Carl Zeiss) operated with AxioVision.

**LC-MS analysis of glucosamine.** Non-labeled myristic acid, non-labeled palmitic acid, [1-$^{13}C_1$]myristic acid (Taiyo Nippon Sanso), and [1-$^{13}C_1$]palmitic acid (Taiyo Nippon Sanso) were neutralized in 200 mM potassium hydroxide to 100 mM final concentration of fatty acids. Approximately 1,200 *R. irregularis* spores were placed in the center hole of 0.75% Phytagel, which was solidified in a 60-mm Petri dish. The Phytagel was then covered with 5 mL of half-strength modified SC medium with one of the following five supplements: 1) 0.5 mM myristic acid, 2) 0.5 mM [1-$^{13}C_1$]myristic acid, 3) 0.5 mM myristic acid and 0.5 mM palmitic acid, 4) 0.5 mM myristic acid and 0.5 mM [1-$^{13}C_1$]palmitic acid, and 5) 0.5 mM [1-$^{13}C_1$]myristic acid and 0.5 mM palmitic acid. The AM fungi were then grown at 28 °C for eight weeks. During the culture period, the liquid medium was changed once a month. After removing the central region containing parent spores, fungal materials including hyphae and myristate-induced secondary spores were recovered from the remaining Phytagel as described above. Glucosamine derived from fungal biomass was extracted according to previous reports (6, 7) with some modifications. Briefly, fungal materials were washed in cold 0.25 M hydrochloric acid solution twice for 5 and 35 min. After centrifugation at 12,000 rpm for 5 min, the fungal pellet was rinsed with Milli-Q water. The sample was then incubated in 1 mL of 0.2 M sodium hydroxide for 6 h at room temperature. After centrifugation, the pellet was further incubated in 1 mL of fresh 0.2 M sodium hydroxide at 70 °C for 17 h. After cooling to room temperature, the sample was washed with Milli-Q water four times and dried at 70 °C for several hours. Acid hydrolysis was performed with 1 mL of 6 M hydrochloric acid solution at 70 °C for 16 h. A total of 40 μL of each extract or glucosamine hydrochloride standard solution (10 μg $mL^{-1}$) was evaporated under reduced pressure, dissolved in 50% acetonitrile, and filtered through 0.45-μm membrane filters. The chromatographic separation was performed using the ACQUITY UPLC System (Waters), where 20 μL of sample was injected onto a COSMOSIL Sugar-D column (4.6 mm × 250 mm, Nacalai Tesque) at 30 °C, and the solute was isocratically eluted using 75% acetonitrile containing 0.05% formic acid as the mobile phase at a flow rate of 0.75 mL $min^{-1}$. A glucosamine standard was detected with a retention time of 2.65 min. The mass spectrometer was operated in a positive ESI mode. The nebulizer and desolvation gas flows were 50 and 700 L $h^{-1}$, respectively. The capillary voltage was set at 2.8 kV, the cone voltage at 45 V, the source temperature at 120 °C, and the desolvation gas temperature at 350 °C. Data were collected using MassLynx version 4.1 (Waters). The relative intensities of the molecular ion peaks of glucosamine ($[M+H]^+$, *m/z* 180.19; and $[M+1+H]^+$, *m/z* 181.19) were monitored. The relative fraction of M+1 with respect to that of M+0 in the glucosamine standard solution was 6.8%.

**$^{13}$C-NMR analysis of triacylglycerol (TAG).** [1-$^{13}C_1$]Myristic acid (Cambridge Isotope Laboratories, Inc.) and non-labeled myristic acid were neutralized in 200 mM potassium hydroxide to a 100 mM final concentration of fatty acids. Further, *R. irregularis* was cultured in modified SC solid medium supplied with 1 mM neutralized non-labeled myristic acid or [1-$^{13}C_1$]myristic acid for 73 days. After removing the central region of the solid medium that contained parent spores, approximately 25 mg and 43 mg of wet fungal materials were recovered respectively from non-labeled myristic acid and [1-$^{13}C_1$]myristic acid-fed plates by gel solubilization as described above followed via vacuum filtration through filter paper. Lipids derived from fungal biomass were extracted with 3 mL of chloroform–methanol (2:1) via sonication for 30 min. After filtration, the lipid extract solutions were concentrated by nitrogen gas. The concentrates were purified by preparative silica gel TLC (Kieselgel 60 $F_{254}$, Merck) using *n*-hexane–diethyl ether–acetic acid (80:30:1) as a developing solvent to yield 1.0 and 1.1 mg of TAG from non-labeled myristic acid and [1-$^{13}C_1$]myristic acid-treated samples, respectively. For NMR analysis, the TAG was dissolved in 600 μL of $CDCl_3$. The $^{13}$C-NMR spectra were recorded at 125 MHz on a JNM-ECZ500R spectrometer (JEOL) using the default parameter settings. The chemical shifts were referenced to the solvent peak ($CDCl_3$ $\delta_C$ 77.0) as an internal standard. 1.6 mg of TAG prepared from 27 mg of wet fungal materials of *R. irregularis* grown in a monoxenic root organ culture (Premier Tech) was also subjected to $^{13}$C-

NMR analysis as above.

**GC-MS analysis of TAG.** *R. irregularis* was incubated for eight weeks in half-strength modified SC medium with one of the following four supplements: 1) 0.5 mM neutralized myristic acid, 2) 0.5 mM neutralized [1-$^{13}C_1$]myristic acid, 3) 0.1 mM potassium myristate and 0.5 mM neutralized palmitic acid, and 4) 0.1 mM potassium myristate and 0.5 mM neutralized [1-$^{13}C_1$]palmitic acid as described in LC-MS analysis of glucosamine. The extraction of lipids from the fungal materials and purification of TAG were performed as described in the $^{13}$C-NMR analysis of TAG. Fatty acid methyl esters were prepared by incubating the purified TAG in 80 µL hexane and 8 µL 1 M methanolic KOH as described previously (8). An aliquot of the upper hexane layer was applied to GC-MS. The GC-MS data were recorded with a GCMS-QP2010 Plus (Shimadzu) and an InertCap 5MS/NP column (25 m×0.25 mm, 0.25 µm film, GL Sciences). Specific conditions were used for chromatographic separation: injection 1 µL (splitless, 60 s valve time), injector temperature 200 °C, carrier gas He (at 0.8 mL min$^{-1}$), transfer line temperature 300 °C, ion source temperature 230 °C, electron energy 70 eV. The temperature of the column oven was programmed in three steps: 60 °C for 2 min, followed by an increase to 160 °C at 25 °C min$^{-1}$, and then an increase to 300 °C at 5 °C min$^{-1}$. The identification and quantification of fatty acid methyl esters were performed in scan mode. The composition of C14:0, C16:0, and C16:1Δ11 fatty acids was determined by calibration with external standards. Positional isomers of C16:1 from AM fungi were determined by comparing the retention times of methyl ester standards of C16:1Δ9 (15.90 min) and C16:1Δ11 (16.07 min). Incorporation rates of [$^{13}C_1$] into TAG were calculated from the peak intensity of molecular ions $[M]^+$ (*m/z* 242, 270, and 268 for C14:0, C16:0, and C16:1Δ11 methyl esters, respectively) and isotopic ions $[M+1]^+$ (*m/z* 243, 271, and 269 for C14:0, C16:0, and C16:1Δ11 methyl esters, respectively) in the electron-impact mass spectra of C14:0, C16:0, and C16:1Δ11 methyl ester peaks equated as follows: [$^{13}C_1$]-incorporation (%) = (($[M+1]^+ - [M]^+ \times$ natural abundance of $[M+1]^+$ isotope) / ($[M]^+ + ([M+1]^+ - [M]^+ \times$ natural abundance of $[M+1]^+$ isotope)) × 100.

**Determination of ATP content.** Five hundred spores of *R. irregularis* were incubated in 100 µL of sterilized water at 28 °C for five days. Potassium myristate was added to the germinating spores at a final concentration of 0.5 mM. For the control, the protonophore carbonyl cyanide *m*-chlorophenylhydrazone (CCCP) was simultaneously added at a final concentration of 50 µM. The germinating spores were incubated for 12 h. After centrifugation at 3,500 rpm for 10 min, 400 µL of phosphate buffered saline (PBS; pH 7.4) was added to the pellet. The spore suspension was transferred to a 2 mL tube containing a metal crusher (TAITEC) and placed on ice. The sample was crushed with a bead crusher (µT-12, TAITEC) at 2,200 rpm for 10 s three times. The crushed fungal materials were transferred to a 1.5 mL tube and centrifuged at

8,000×*g* for 2 min at 4 °C. The supernatant was recovered and used for determination of ATP and protein concentration. ATP concentration was measured using the CellTiter-Glo Luminescent Cell Viability Assay kit (Promega) following the manufacturer's instructions. Protein concentration was assayed using the Qubit Protein Assay Kits (Thermo Fisher Scientific). ATP content in the germinating spores was calculated in nmol $mg^{-1}$ of protein.

**Quantitative RT-PCR.** *R. irregularis* was grown in an immobilized cell culture system with modified SC medium containing 0.5 mM potassium myristate without sugars for three weeks. Subsequently, Phytagel tablets containing fungal materials were incubated in a modified SC medium without fatty acids for 11 days to induce fatty acid starvation. During the first three days of starvation, the culture medium was exchanged every day. After starvation, 100 mM potassium myristate to a final concentration of 0.5 mM was added to half of the samples, and the same amount of sterilized water was added to the remaining half of the samples. After a 3-h incubation, fungal hyphae protruding outside a Phytagel tablet were recovered using forceps and immediately immersed in 500 µL of RNAiso Plus (Takara Bio). Contamination by phytagel resulted in a reduced RNA yield. The fungal hyphae were crushed using the bead crusher µT-12 with a metal crusher at 2,200 rpm for 10 s three times with cooling on ice. RNA was extracted using RNAiso Plus following the manufacturer's instructions. Isolated RNA was treated with a TURBO DNA-free Kit (Thermo Fisher Scientific) to eliminate contaminating genomic DNA. cDNA was synthesized using a ReverTra Ace qPCR RT Kit (TOYOBO). Semiquantitative PCR was conducted using the StepOne Real-Time PCR System (Thermo Fisher Scientific) with a THUNDERBIRD SYBR qPCR Mix (TOYOBO). Primers used for the qRT-PCR are shown in *SI Appendix*, Table S4. Melting curve analysis confirmed the single peaks. Furthermore, *R. irregularis* elongation factor 1 beta (*EF-1β*) and actin (*Act*) genes were used for normalization of gene expression levels of target genes. For each genotype and treatment five to six biological replicates were tested and each sample was represented by two to three technical replicates.

**Statistical analysis.** All statistical analyses were performed using R version 3.5.2. Levene's tests were applied to check for heteroscedasticity between treatment groups. Data were transformed as $\log_{10}(x + 0.5)$ where necessary. To examine the differences among experimental groups, data were analyzed with Student's *t*-test, Tukey's HSD test, Wilcoxon–Mann–Whitney test, and Steel–Dwass test, as appropriate. Differences at $P < 0.05$ were considered significant.

**References**

1. Y. Kobayashi, T. Maeda, K. Yamaguchi, H. Kameoka, S. Tanaka, T. Ezawa, S. Shigenobu, M. Kawaguchi, The genome of *Rhizophagus clarus* HR1 reveals a common genetic basis for auxotrophy among arbuscular mycorrhizal fungi. *BMC Genomics* **19,**

465 (2018).
2. F. A. de Souza, G. A. Kowalchuk, P. Leeflang, J. A. van Veen, E. Smit, PCR-denaturing gradient gel electrophoresis profiling of inter- and intraspecies 18S rRNA gene sequence heterogeneity is an accurate and sensitive method to assess species diversity of arbuscular mycorrhizal fungi of the genus *Gigaspora*. *Appl. Environ. Microbiol.* **70,** 1413–1424 (2004).
3. S. Cranenbrouck, L. Voets, C. Bivort, L. Renard, D.-G. Strullu, S. Declerck, "Methodologies for in vitro cultivation of arbuscular mycorrhizal fungi with root organs" in *In Vitro Culture of Mycorrhizas*, S. Declerck, D.-G. Strullu, J. A. Fortin, Eds. (Springer, 2005), pp. 342–375.
4. H. Kameoka, I. Tsutsui, K. Saito, Y. Kikuchi, Y. Handa, T. Ezawa, H. Hayashi, M. Kawaguchi, K. Akiyama, Stimulation of asymbiotic sporulation in arbuscular mycorrhizal fungi by fatty acids. *Nature Microbiol.* **4,** 1654–1660 (2019).
5. J. M. Phillips, D. S. Hayman, Improved procedures for clearing roots and staining parasitic and vesicular-arbuscular mycorrhizal fungi for rapid assessment of infection. *Trans. Br. Mycol. Soc.* **55**, 158–161 (1970).
6. A. Ekblad A, T. Näsholm, Determination of chitin in fungi and mycorrhizal roots by an improved HPLC analysis of glucosamine. *Plant Soil* **178**, 29–35 (1996).
7. M. A. Olofsson, D. Bylund, Liquid chromatography with electrospray ionization and tandem mass spectrometry applied in the quantitative analysis of chitin-derived glucosamine for a rapid estimation of fungal biomass in soil. *Int. J. Anal. Chem.* **2016**, 8 (2016).
8. K. Ichihara, A. Shibahara, K. Yamamoto, T. Nakayama, An improved method for rapid analysis of the fatty acids of glycerolipids. *Lipids* **31,** 535–539 (1996).

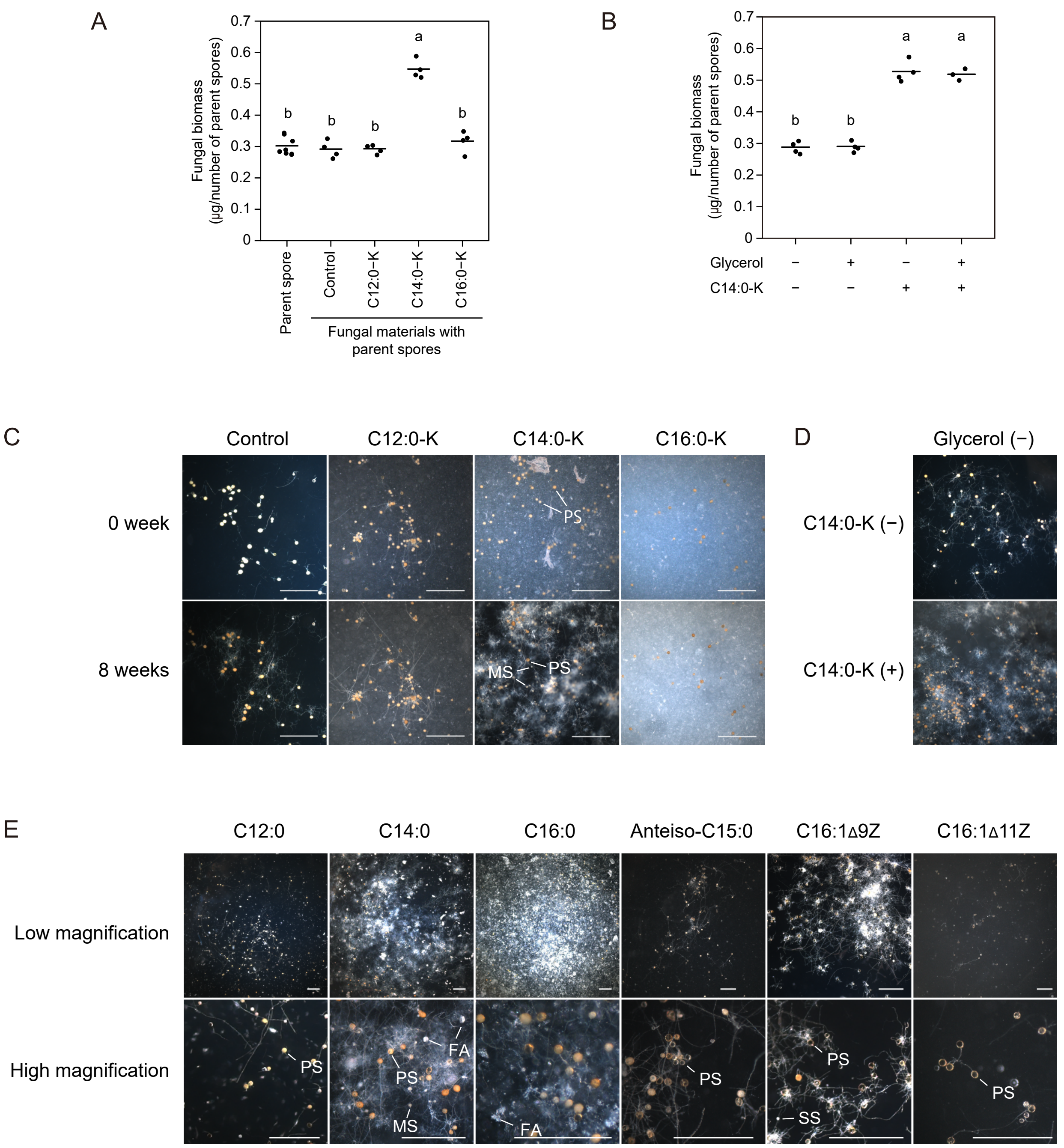


**Fig. S1.** *R. irregularis* cultured in the solid media supplemented with fatty acids. (*A*) Biomass of parent spores ($n$ = 8) and fungal materials cultured in the modified SC solid medium supplemented with potassium salts of fatty acids after eight weeks of cutivation ($n$ = 4). (*B*) Biomass of fungal materials cultured in the presence or absence of 1 mM glycerol and 0.5 mM potassium myristate (C14:0-K) after eight weeks of cutivation ($n$ = 3–4). Fungal biomass was calculated by dividing the total fungal dry weight in each well by the number of parent spores. Horizontal lines show mean values. The same lowercase letter indicates no significant difference (Tukey’ s test, $P < 0.05$). Fungal growth in modified SC medium supplemented with 1 mM potassium salts of fatty acids (*C*) and 1 mM fatty acids (*E*) after eight weeks of cultivation. (*D*) Fungal growth in modified SC medium without glycerol after eight weeks of cultivation. See *SI Appendix*,Table S2 for sample details. FA, fatty acid precipitate; MS, myristate-induced spore; PS, parent spore; and SS, secondary spore. Scale bars: 1 mm.

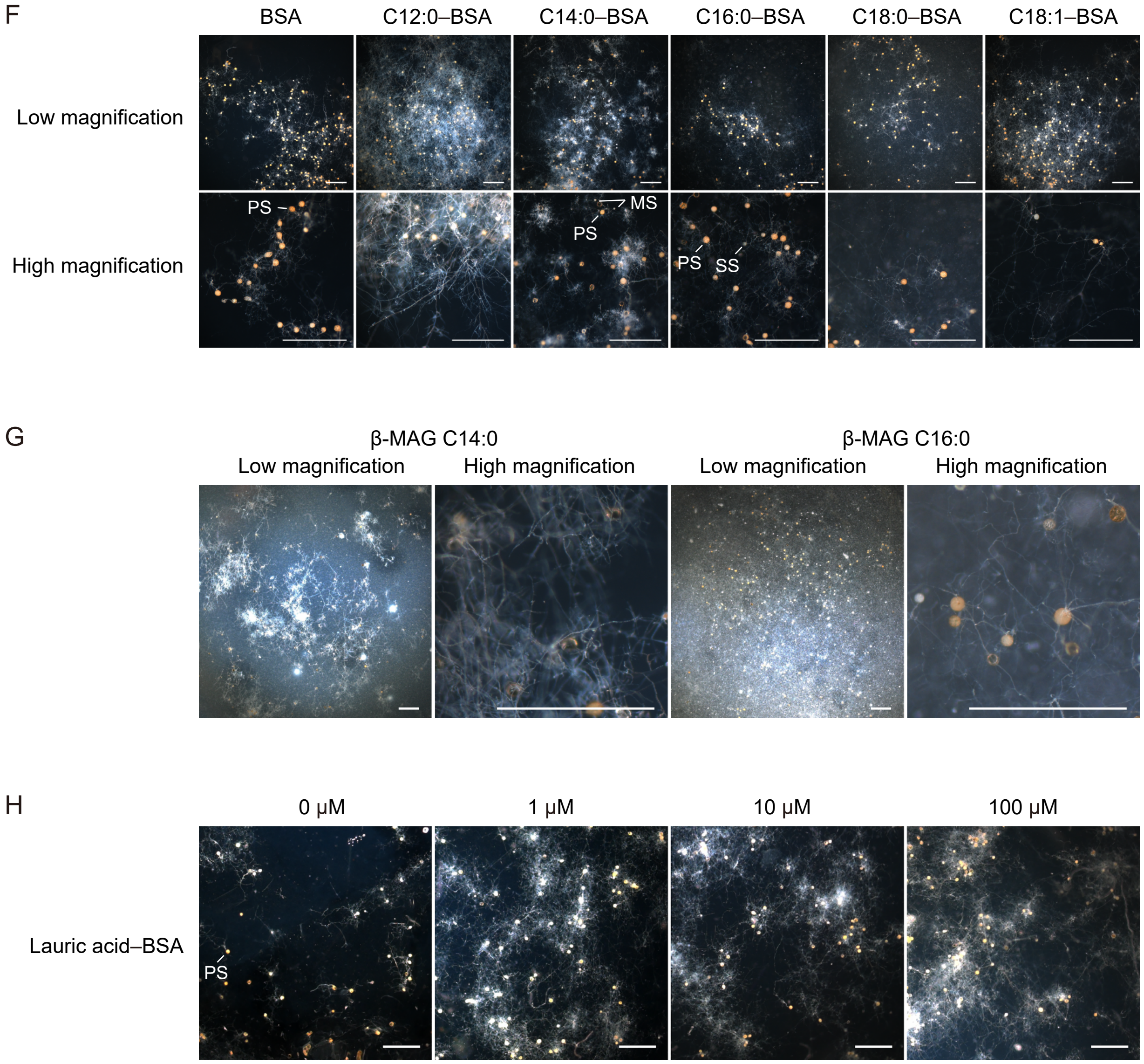


**Fig. S1.** continued. Fungal growth in modified SC medium supplemented with 0.5 mM fatty acid–BSA conjugates (*F*) and 1 mM *sn*-2 monoacylglycerols (β-MAGs) (*G*) after eight weeks of cultivation. (*H*) Effect of different concentrations of lauric acid (C12:0)–BSA conjugates on fungal growth after eight weeks of cultivation. MS, myristate-induced spore; PS, parent spore; and SS, secondary spore. Scale bars: 1 mm.

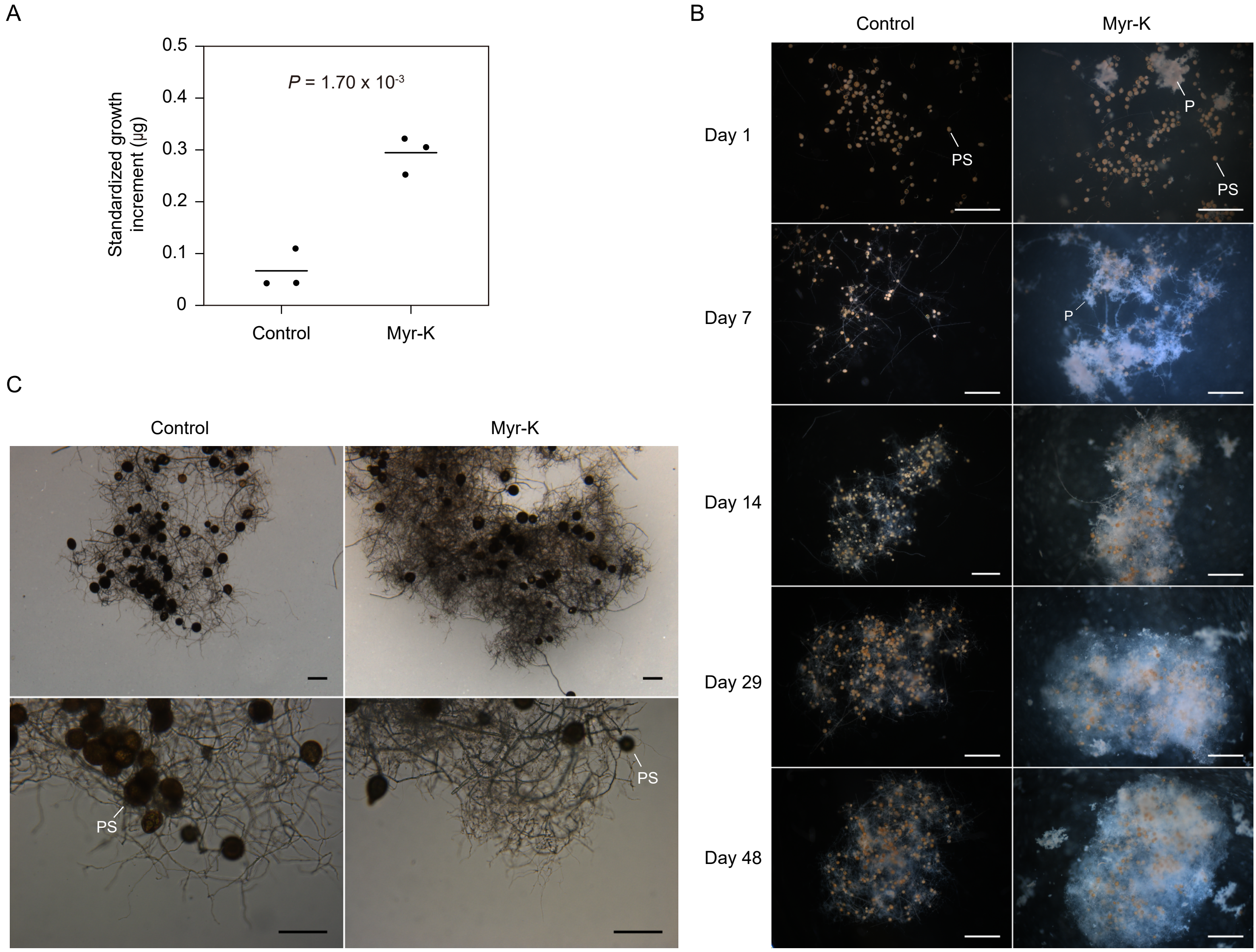


**Fig. S2.** Liquid culture of *R. irregularis*. (*A*) Biomass production in the modified SC liquid medium supplemented with or without 0.5 mM potassium myristate (Myr-K) after eight weeks of cultivation. Horizontal lines indicate mean values ($n$ = 3). The $P$ value is based on the Student' s $t$-test. (*B*) Time course of fungal growth in the liquid medium without fatty acids or with potassium myristate. When potassium myristate was added to the medium, precipitates of the metal soaps, which attached to the hyphal surface, were apparent. (*C*) Fungal growth in the liquid medium after eight weeks of cultivation. To remove the metal soaps attached to the hyphal surface, fungal hyphae were washed with citrate buffer including 1% Triton X-100. Images were captured using an inverted microscope. Lower panels are magnified images of fungal hyphae. See *SI Appendix*,Table S2 for sample details. P, precipitate of metal soap; and PS, parent spore. Scale bars: 1,000 μm (*B*) and 200 μm (*C*).

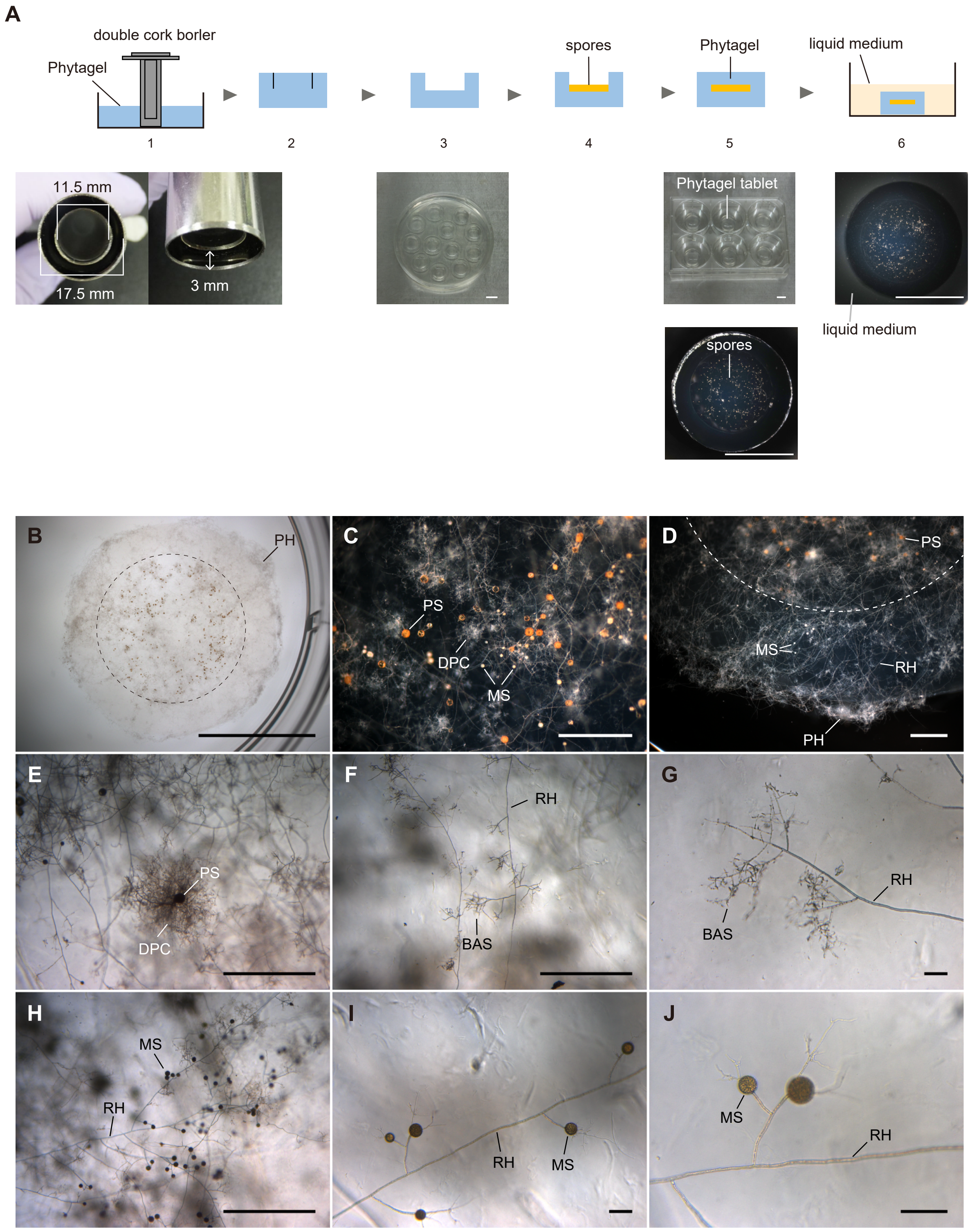


**Fig. S3.** Immobilized cell culture of *R. irregularis*. (*A*) Schematic presentation of the immobilized cell culture system. Step 1 and 2: Phytagel tablets with a circular incision were cut out using a sterile double cork borer. Step 3: The gel within the circular incision on the top side of the gel tablet was removed. Step 4 and 5: Parent spores were placed in the hole and covered with Phytagel. Step 6: The Phytagel tablets, which contained spores, were transferred into a six-well culture plate. Each well was filled with the modified SC liquid medium with fatty acids. (*B*) Fungal growth by immobilized cell culture containing half-strength modified SC medium supplemented with 0.5 mM potassium myristate after eight weeks of cultivation. A dotted circle shows the area in which parent spores are placed. Magnified images of (*C*) the central region and (*D*) peripheral area of a Phytagel tablet. (*E*) DPC-like structures formed around the parent spore. (*F*) BAS formed along the runner hyphae. (*G*) Magnified image of BAS. (*H–J*) Myristate-induced secondary spores formed along the runner hyphae. (*I* and *J*) Magnified images of myristate-induced spores. See *SI Appendix*,Table S2 for sample details. BAS, branched absorbing structure; DPC, densely packed coil; MS, myristate-induced spore; PH, hypha protruding outside a Phytagel tablet; PS, parent spore; and RH, runner hypha. Scale bars: 10 mm (*A* and *B*), 1 mm (*C–F* and *H*), and 0.1 mm (*G*, *I*, and *J*).

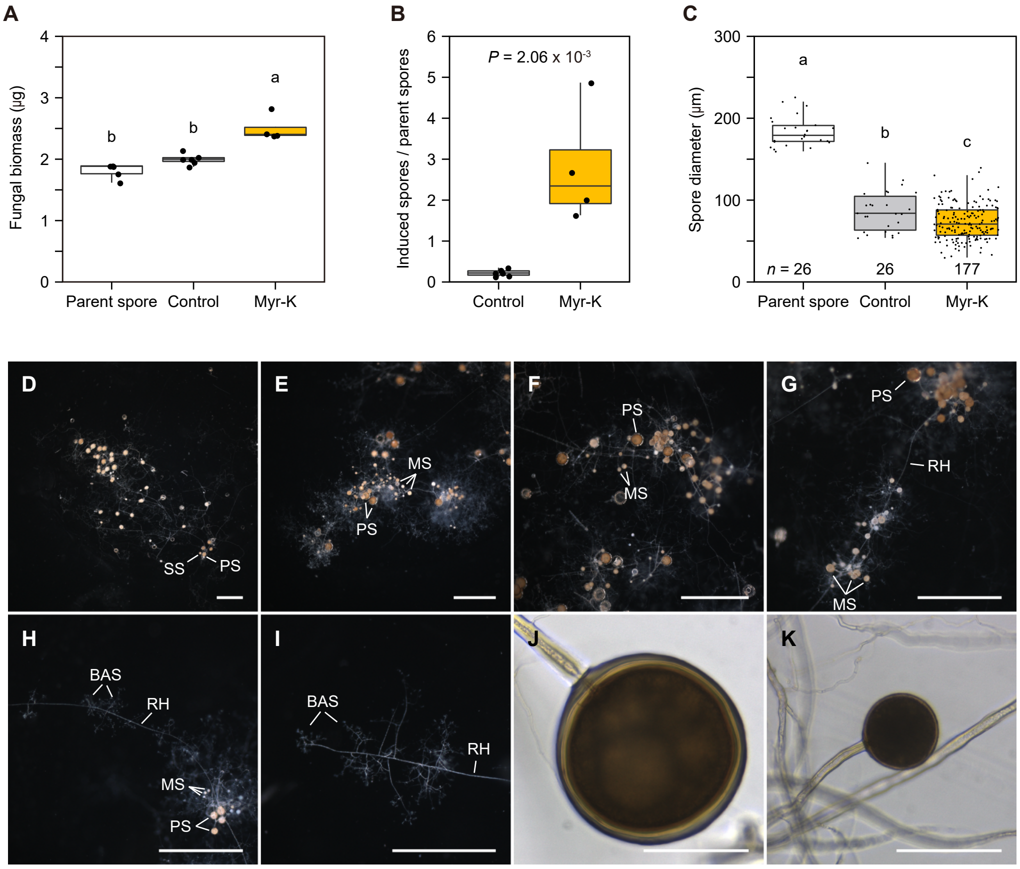


**Fig. S4.** Asymbiotic culture of *Rhizophagus clarus*. (*A*) Biomass of parent spores and fungal materials in the immobilized culture system with or without potassium myristate (Myr-K) after 12 weeks of cultivation ($n$ = 4–6). Fungal biomass was calculated by dividing the total fungal dry weight in each well by the number of parent spores. Number (*B*) and diameter (*C*) of *R. clarus* spores. See *SI Appendix*, Table S2 for sample details. For each boxplot, the boxes show the first quartile, the median, and the third quartile; the whiskers reach to the 1.5× interquartile range, and data points for each treatment are displayed. The same lowercase letter indicates no significant difference (Tukey' s test, $P < 0.05$ (*A* and *C*)). The *P* value in (*B*) is based on the Student' s *t*-test ($n$ = 4–6). Fungal growth without fatty acids (*D*) or with 0.5 mM potassium myristate (*E*–*K*) for 12 weeks. (*D*) *R. clarus* in the absence of fatty acids produced several germ tubes from which a few secondary spores were generated. (*E*) Extensive hyphal branching and secondary spore formation around parent spores in the presence of myristate. (*F*) Myristate-induced secondary spores generated around the parent spores. (*G*) Myristate-induced spores formed along the runner hyphae. (*H*) BAS-like structures along runner hyphae. Magnified images of BAS (*I*), a parent spore (*J*), and a myristate-induced secondary spore (*K*). BAS, branched absorbing structure; MS, myristate-induced spore; PS, parent spore; RH, runner hypha; and SS, secondary spore. Scale bars: 1 mm (*D*–*I*) and 100 µm (*J* and *K*).

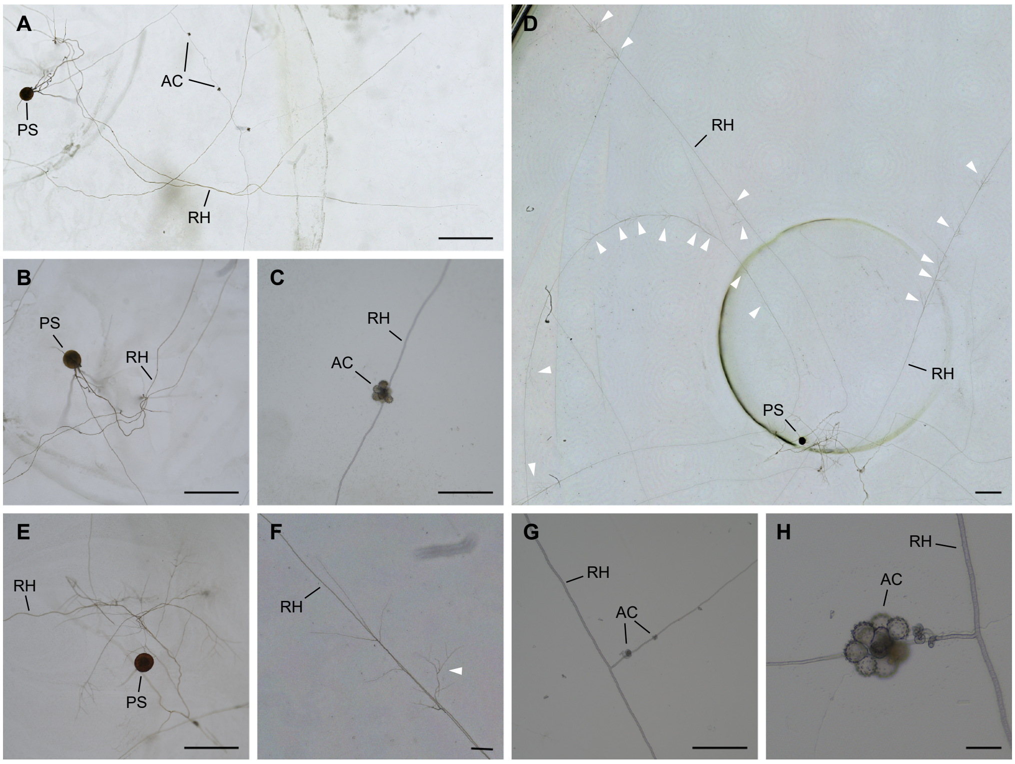


**Fig. S5.** Asymbiotic culture of *Gigaspora margarita*. AM fungi were cultured in the immobilized cell culture system without fatty acids (*A*–*C*) or with 0.5 mM potassium myristate (*D*–*H*) for 12 weeks. (*A*) A germinating spore of *G. margarita* produced several runner hyphae along which a few auxiliary cells were formed. Magnified images of a germinating spore (*B*) and an auxiliary cell (*C*). (*D*) *G. margarita* in the presence of myristate produced long runner hyphae from which BAS-like structures (arrowheads) were frequently generated. (*E*) Branching of runner hyphae around a parent spore. (*F*) Magnified image of BAS (arrowhead) stemmed from a runner hypha. (*G*) Auxiliary cells. (*H*) Magnified image of an auxiliary cell. AC, auxiliary cell; PS, parent spore; and RH, runner hypha. Scale bars: 1 mm (*A*, *B*, *D*, and *E*), 200 μm (*C*, *F*, and *G*), and 50 μm (*H*).

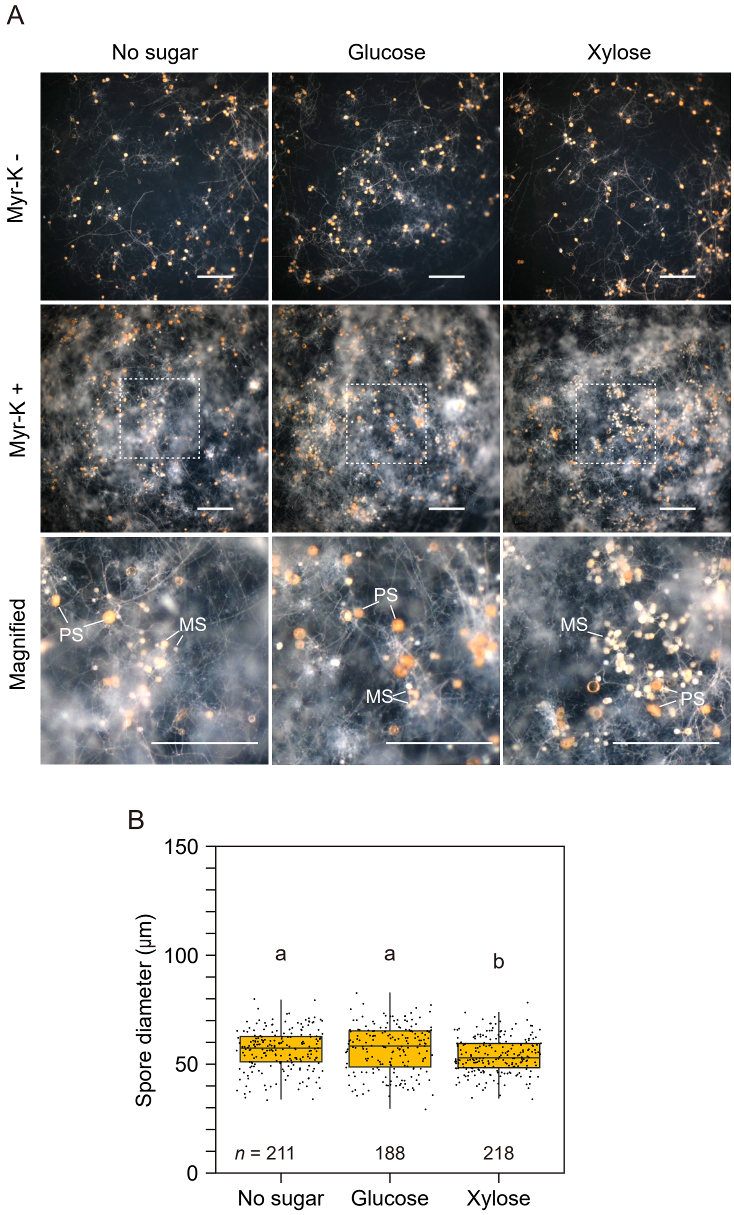


**Fig. S6.** Effects of sugars in combination with myristate on the growth and sporulation of *R. irregularis*. (*A*) Fungal growth in the immobilized cell culture containing half-strength modified SC medium supplemented with combinations of 0.5 mM potassium myristate (Myr-K) and 5 mM sugars after two months of cultivation. Images were taken by a dissecting microscope. Lower panels show the magnified images of dotted areas in the middle panels. Yellow-orange spores are parent spores (PS) and small spores in white to light-yellow color are myristate-induced spores (MS). Scale bars: 1 mm. (*B*) Size of the myristate-induced spores in immobilized cell culture system. Boxes show the first quartile, the median, and the third quartile; the whiskers reach to the 1.5× interquartile range, and data points for each treatment are displayed. The same lowercase letter indicates no significant difference (Steel–Dwass test, $P < 0.05$).

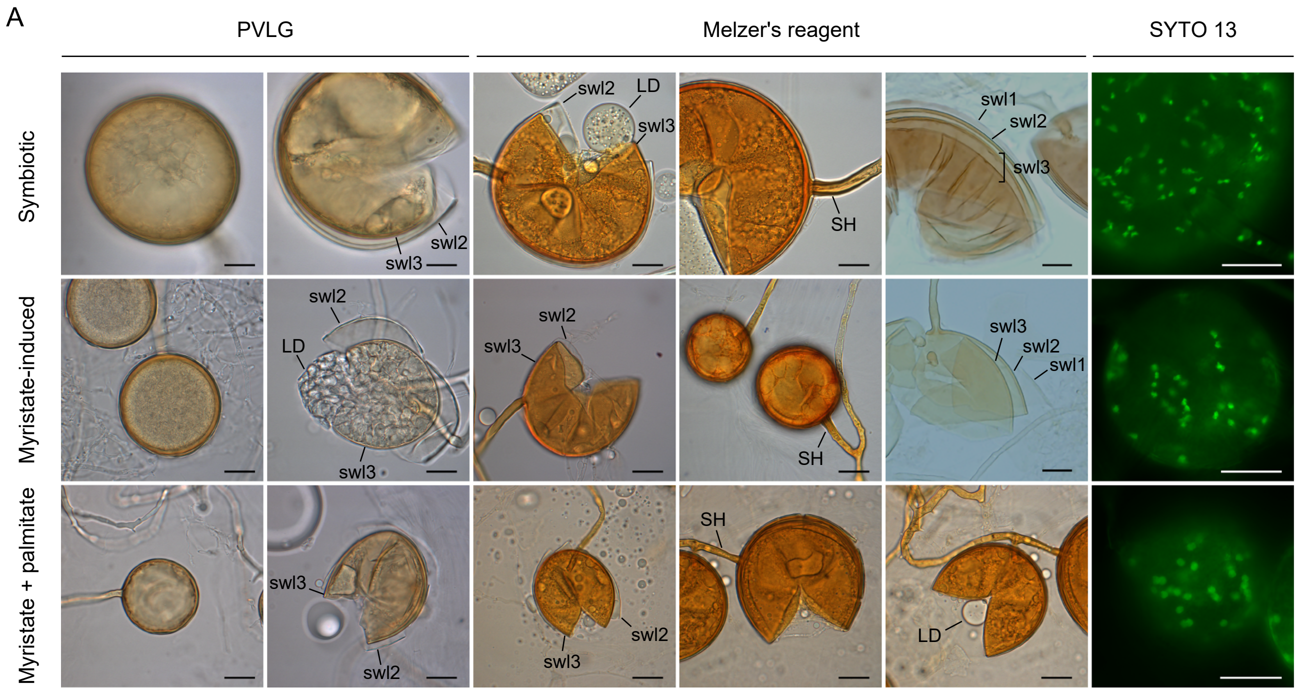


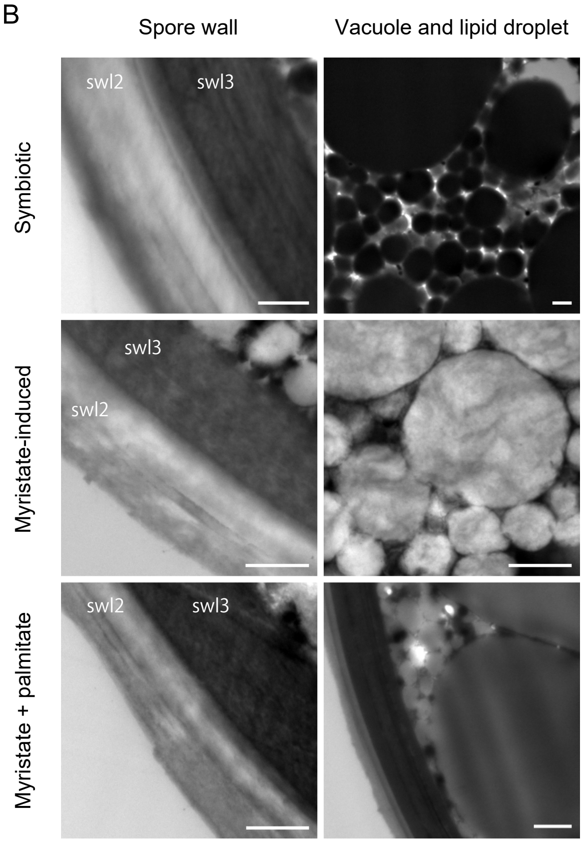


**Fig. S7.** Spore morphology. (*A*) Symbiotically generated spores and myristate-induced spores of *R. irregularis* that were produced using an immobilized cell culture system supplemented with 0.5 mM potassium myristate or 0.1 mM potassium myristate plus 0.5 mM potassium palmitate. Intact or crushed spores were mounted in PVLG or Melzer' s reagent. Spores appear yellow-brown in color and have apparent two spore wall layers (swl2 and swl3). The outmost spore wall layer (swl1) that naturally sloughs off was occasionally observed. Subtending hyphae of all type of spores exhibit a cylindrical shape. Myristate-induced spores were smaller than symbiotically generated spores. Lipid droplets in symbiotically generated spores and myristate-induced spores that were produced in palmitate-supplemented culture media were in liquid-like state, whereas myristate-induced spores cultured in the absence of palmitate had solid-like state. Nuclei in the spores were stained with SYTO 13 Green Fluorescent Nucleic Acid Stain and observed by epifluorescence microscopy. Many nuclei were observed in both myristate-induced and symbiotically generated spores. LD, lipid droplet; and SH, subtending hypha. (*B*) Transmission electron micrographs of the spore wall and the inside of the spores. Spore wall comprises two prominent layers (swl2 and swl3). Spore wall layers (swl2 and swl3) of myristate-induced spores were less thick than those of symbiotically generated spores. Many vacuoles and lipid droplets were visible in all type of spores. Scale bars: 20 μm (*A*) and 1 μm (*B*).

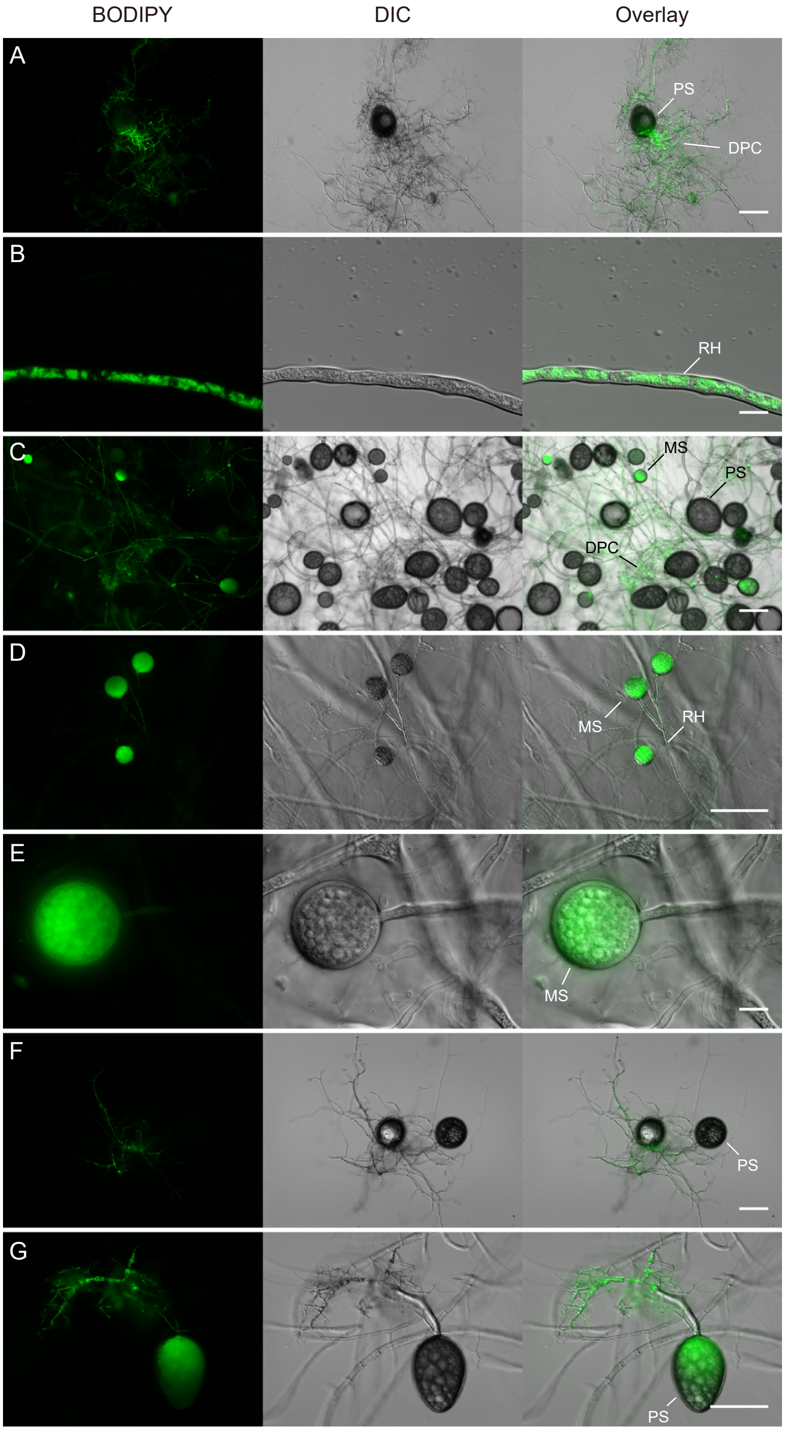


**Fig. S8.** Uptake of the fluorescently labelled fatty acid derivative $C_1$-BODIPY 500/510 $C_{12}$ by *R. irregularis*. (*A*–*E*) Fluorescent images of $C_1$-BODIPY 500/510 $C_{12}$ and superimposed differential interference contrast (DIC) images of AM fungal materials cultivated using an immobilized cell culture system. AM fungi were incubated in the modified SC medium containing 0.5 mM potassium myristate for six weeks and then stained with the fluorescent probe for one day (*A*, *C*, and *D*) or five days (*B* and *E*). (*A*) Parent spore and DPCs. (*B*) Runner hypha. (*C*) Parent spores and myristate-induced secondary spores. (*D*) Secondary spores formed along runner hypha. (*E*) Secondary spore. (*F* and *G*) Fluorescent images of germinating spores. Seven-day-old germinating spores incubated in sterile water were stained with the fluorescent probe for 10 min (*F*) or seven days (*G*). DPC, densely packed coil; MS, myristate-induced spore; PS: parent spore; and RH, runner hypha. Scale bars: 100 µm (*A*, *C*, *D*, *F*, and *G*) and 10 µm (*B* and *E*).

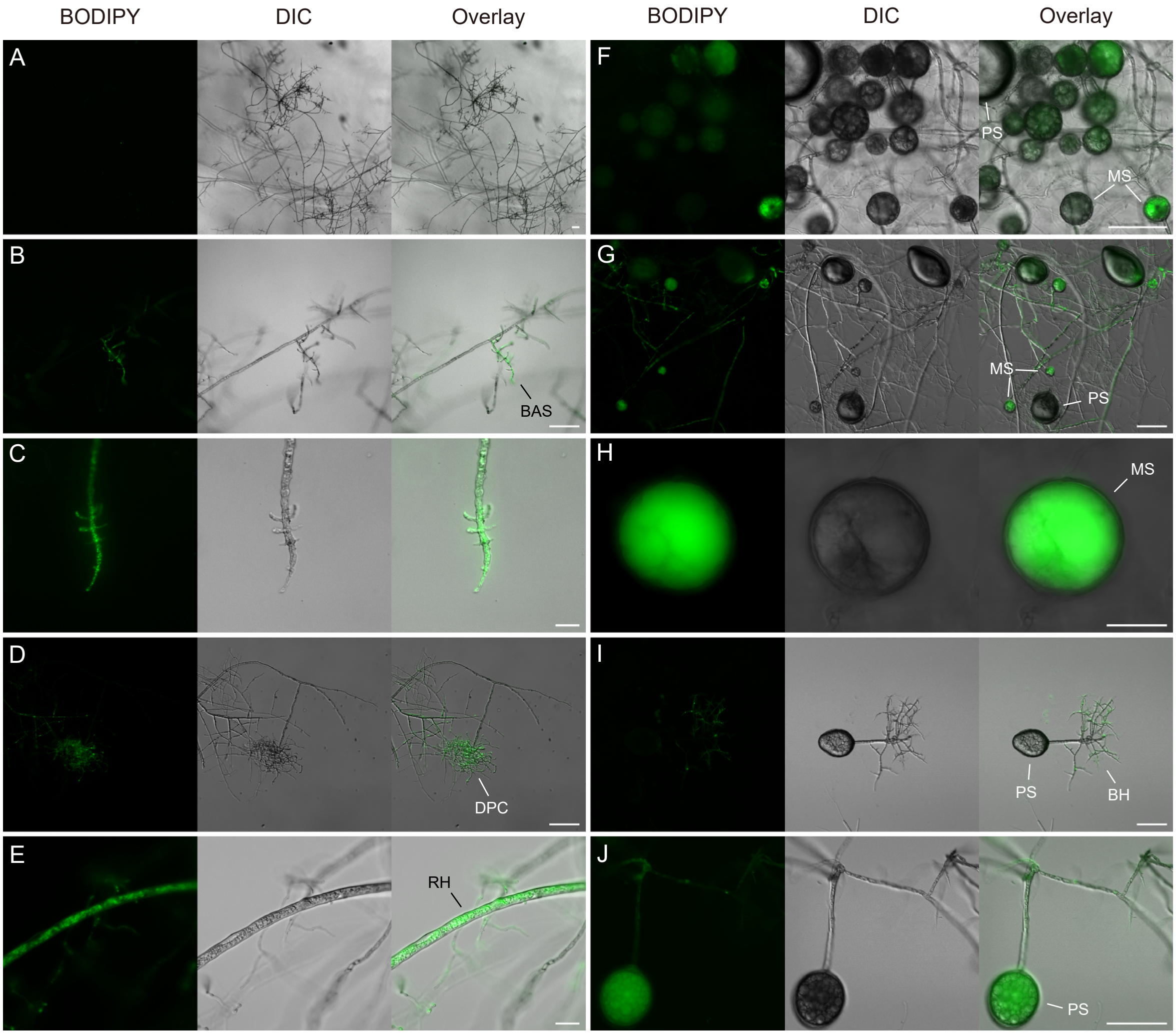


**Fig. S9.** Uptake of the fluorescently labelled fatty acid derivative BODIPY FL $C_{16}$ by *R. irregularis*. (*A*–*H*) Fluorescent images of BODIPY FL $C_{16}$ and superimposed differential interference contrast (DIC) images of AM fungal materials, which were cultivated using an immobilized cell culture system. AM fungi were incubated in the modified SC medium containing 0.5 mM potassium myristate for six weeks and then stained with the fluorescent probe for 10 min (*A*), 4 h (*B* and *C*), three days (*D*–*F*), or nine days (*G* and *H*). (*A* and *B*) BAS. (*C*) Hyphal tip stemmed from BAS. (*D*) DPC. (*E*) Runner hypha. (*F* and *G*) Parent spores and myristate-induced secondary spores. (*H*) Secondary spore. (*I* and *J*) Fluorescent images of germinating spores. Seven-day-old germinating spores incubated in sterile water were stained with the fluorescent probe for 10 min (*I*) or seven days (*J*). BAS, branched absorbing structure; BH, branching hypha; DPC, densely packed coil; MS, myristate-induced spore; PS: parent spore; and RH, runner hypha. Scale bars: 100 μm (*A*, *B*, *D*, *F*, *G*, *I*, and *J*) and 20 μm (*C*, *E*, and *H*).

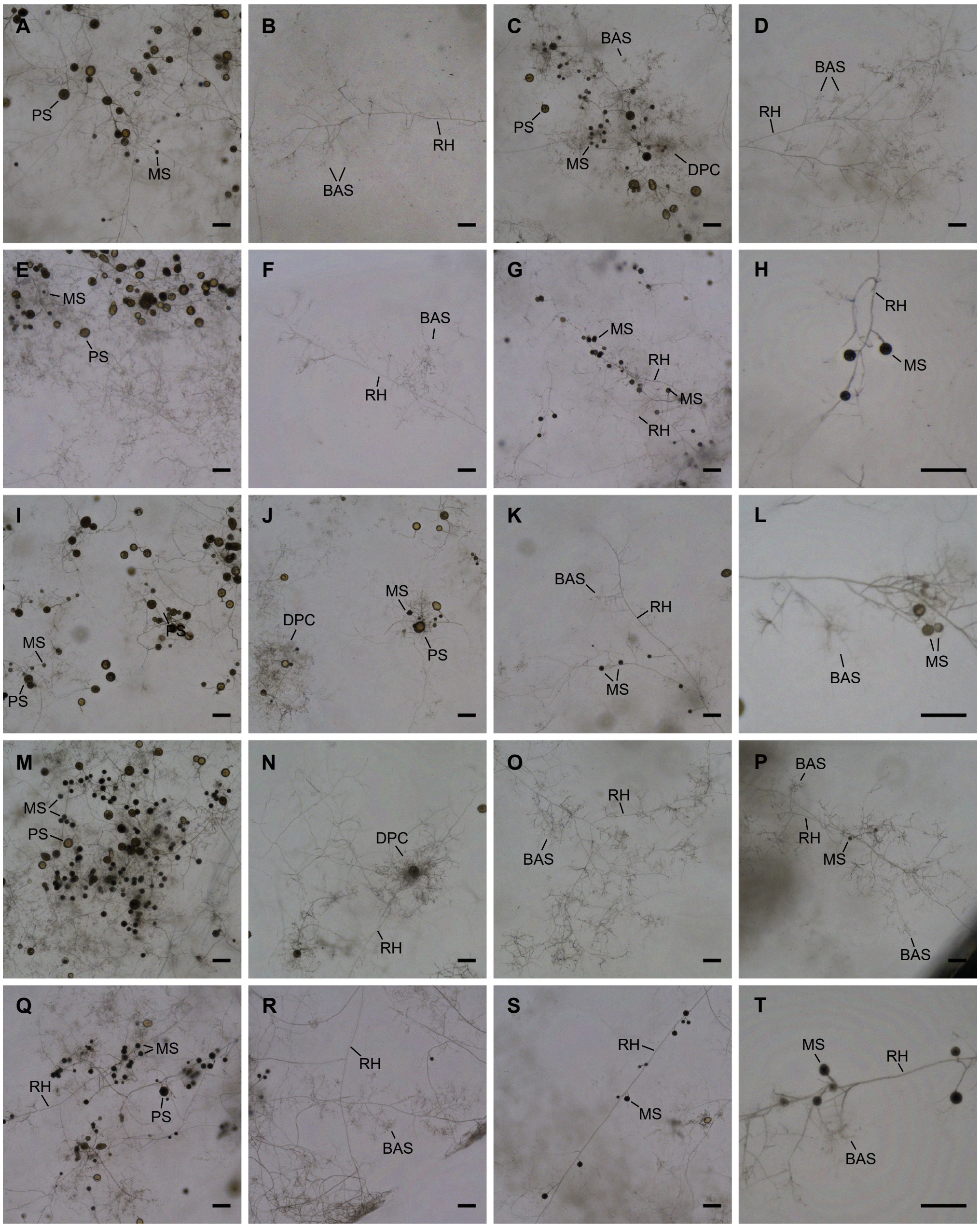


**Fig. S10.** *R. irregularis* growth in the presence of mixtures of fatty acids. AM fungi were cultured in the immobilized cell culture system for eight weeks. The medium contained 0.1 mM potassium myristate (*A* and *B*), 0.5 mM potassium myristate (*C* and *D*), 0.1 mM potassium myristate with 0.5 mM C16:0 *sn*-2 monoacylglycerols (*E*–*H*), 0.1 mM potassium myristate with 0.5 mM potassium palmitate (*I*–*L*), 0.5 mM potassium myristate with 0.5 mM C16:0 *sn*-2 monoacylglycerols (*M*–*P*), or 0.5 mM potassium myristate with 0.5 mM potassium palmitate (*Q*–*T*). (*A*, *C*, *E*, *I*, *M*, and *Q*) Myristate-induced secondary spores generated around the parent spores. (*G*, *H*, *K*, *L*, *P*, *S*, and *T*) Myristate-induced spores formed along the runner hyphae. (*B*, *D*, *F*, *K*, *O*, *P*, and *R*) Branching of runner hyphae and the formation of BAS. (*J* and *N*) DPC-like structures formed around a parent spore. BAS, branched absorbing structure; DPC, densely packed coil; MS, myristate-induced spore; PS, parent spore; and RH, runner hypha. Scale bars: 200 μm.

**Table S1.** Composition of the modified SC medium without fatty acids and sugars.

| Chemicals | mg $L^{-1}$ | Chemicals | mg $L^{-1}$ |
|---|---|---|---|
| $(NH_4)_2SO_4$ | 660 | Alanine | 38 |
| $KH_2PO_4$ | 1000 | Arginine, HCl | 38 |
| $MgSO_4$ | 500 | Asparagine | 38 |
| NaCl | 100 | Aspartic acid | 38 |
| $CaCl_2$ | 100 | Cysteine | 38 |
| Boric acid | 0.5 | Glutamine | 38 |
| $CuSO_4$ | 0.04 | Glutamic acid | 38 |
| KI | 0.1 | Glycine | 38 |
| $FeCl_3$ | 0.2 | Histidine | 38 |
| $MnSO_4$ | 0.4 | Isoleucine | 38 |
| $Na_2MoO_5$ | 0.2 | Leucine | 190 |
| $ZnSO_4$ | 0.4 | Lysine | 38 |
| Glycerol | 92 | Methionine | 38 |
| Biotin | 0.02 | Phenylalanine | 38 |
| Ca pantothenate | 0.4 | Proline | 38 |
| Folic acid | 0.002 | Serine | 38 |
| Inositol | 40 | Threonine | 38 |
| Niacin | 5.4 | Tryptophan | 38 |
| Para-aminobenzoic acid | 4.2 | Tyrosine | 38 |
| Pyridoxine, HCl | 0.4 | Valine | 38 |
| Pyridoxal phosphate | 5 | Phytagel | 3000 |
| Riboflavin | 0.2 | | |
| Thiamine, HCl | 5.4 | | |
| Adenine sulfate | 9 | | |
| Uracil | 38 | | |

**Table S2.** Samples used in the present study.

[a]Immobilized cell culture, [b]*in vitro* culture using carrot hairy roots, [c]weeks after cultivation, [d]C, confocal laser scanning microscope; D, dissecting microscope; E, epifluorescence microscope; F, fully focused composite image; I, inverted microscope; IK, inverted microscope of Keyence; L, light microscope; and T, transmission electron microscope.

| Figure | Culture | Medium | Fatty acid | Sugar | Week[c] | Replication | Microscopy[d] | Scale (µm) |
|---|---|---|---|---|---|---|---|---|
| Fig. 1A | Solid | SC | None, Control | None | 8 | 4 | - | - |
| | Solid | SC | 0.05% BSA, BSA | None | 8 | 3 | - | - |
| | Solid | SC | 1 mM pottasium laurate, C12:0-K | None | 8 | 4 | - | - |
| | Solid | SC | 1 mM lauric acid, C12:0 | None | 8 | 4 | - | - |
| | Solid | SC | 0.5 mM lauric acid-BSA, C12:0-BSA | None | 8 | 4 | - | - |
| | Solid | SC | 1 mM pottasium myristate, C14:0-K (Myr-K) | None | 8 | 4 | - | - |
| | Solid | SC | 1 mM myristic acid, C14:0 | None | 8 | 4 | - | - |
| | Solid | SC | 0.5 mM myristic acid-BSA, C14:0-BSA | None | 8 | 4 | - | - |
| | Solid | SC | 1 mM pottasium palmitate, C16:0-K (Pal-K) | None | 8 | 4 | - | - |
| | Solid | SC | 1 mM palmitic acid, C16:0 | None | 8 | 3 | - | - |
| | Solid | SC | 0.5 mM palmitic acid-BSA, C16:0-BSA | None | 8 | 4 | - | - |
| | Solid | SC | 1 mM 12-methyltetradecanonic acid, Antesio-C15:0 | None | 8 | 4 | - | - |
| | Solid | SC | 1 mM palmitoleic acid, C16:1Δ9Z | None | 8 | 3 | - | - |
| | Solid | SC | 1 mM palmitvaccenic acid, C16:1Δ11Z | None | 8 | 4 | - | - |
| | Solid | SC | 0.5 mM stearic acid-BSA, C18:0-BSA | None | 8 | 4 | - | - |
| | Solid | SC | 0.5 mM oleic acid-BSA, C18:1-BSA | None | 8 | 4 | - | - |
| | Solid | SC | 1 mM C14:0 *sn* -2 monoacylglycerol, β-MAG C14:0 | None | 8 | 4 | - | - |
| | Solid | SC | 1 mM C16:0 *sn* -2 monoacylglycerol, β-MAG C16:0 | None | 8 | 4 | - | - |
| Fig. 1B | Solid | SC | None | None | 2, 4, 6, 8 | 6 | - | - |
| | Solid | SC | 0.2 mM Myr-K | None | 2, 4, 6, 8 | 5-6 | - | - |
| | Solid | SC | 0.5 mM Myr-K | None | 2, 4, 6, 8 | 5-6 | - | - |
| | Solid | SC | 1.0 mM Myr-K | None | 2, 4, 6, 8 | 5-6 | - | - |
| Fig. 1C | ICC[a] | 0.5× SC | None | None | 8 | 6 | - | - |
| | ICC | 0.5× SC | 0.5 mM Myr-K | None | 8 | 6 | - | - |
| | ICC | 0.5× SC | None | 5 mM glucose | 8 | 6 | - | - |
| | ICC | 0.5× SC | 0.5 mM Myr-K | 5 mM glucose | 8 | 6 | - | - |
| | ICC | 0.5× SC | None | 5 mM xylose | 8 | 6 | - | - |
| | ICC | 0.5× SC | 0.5 mM Myr-K | 5 mM xylose | 8 | 6 | - | - |
| Fig. 1D | Solid | SC | None | None | 2 | - | I | 200 |
| Fig. 1E | Solid | SC | None | None | 8 | - | D | 200 |
| Fig. 1F | Solid | SC | 1 mM Myr-K | None | 8 | - | D | 200 |
| Fig. 1G | Solid | SC | 1 mM Myr-K | None | 2 | - | I | 200 |
| Fig. 1H | Solid | SC | 1 mM Myr-K | None | 8 | - | D | 1,000 |
| Fig. 1I | Solid | SC | 1 mM Myr-K | None | 5 | - | I, F | 200 |
| Fig. 1J | Solid | SC | 1 mM Myr-K | None | 5 | - | I, F | 200 |
| Fig. 1K | Solid | SC | 1 mM Myr-K | None | 5 | - | I, F | 200 |
| Fig. 1L | Solid | SC | 1 mM Myr-K | None | 5 | - | I, F | 200 |
| Fig. 1M | Solid | SC | 1 mM Myr-K | None | 5 | - | I, F | 200 |
| Fig. 1N | Solid | SC | 1 mM Myr-K | None | 8 | - | D | 200 |
| Fig. 1O | Solid | SC | 1 mM Myr-K | None | 5 | - | I | 200 |
| Fig. 1P | Solid | SC | 1 mM Myr-K | None | 5 | - | I | 200 |
| Fig. 1Q | ICC | 0.5× SC | None | None | 8 | - | D | 1,000 |
| Fig. 1R | ICC | 0.5× SC | 0.5 mM Myr-K | None | 8 | - | D | 1,000 |
| Fig. 2A | Solid | SC | None | None | 2, 4, 6, 8 | 3 | - | - |
| | Solid | SC | 0.2 mM Myr-K | None | 2, 4, 6, 8 | 5-6 | - | - |
| | Solid | SC | 0.5 mM Myr-K | None | 2, 4, 6, 8 | 5-6 | - | - |
| Fig. 2B | Solid | SC | None | None | 8 | 4 | - | - |
| | Solid | SC | 0.2 mM Myr-K | None | 8 | 205 | - | - |
| | Solid | SC | 0.5 mM Myr-K | None | 8 | 268 | - | - |
| | - | - | None | None | 0 | 206 | - | - |
| Fig. 2C | ICC | 0.5× SC | 0.5 mM Myr-K | None | 8 | 6 | - | - |
| | ICC | 0.5× SC | 0.5 mM Myr-K | 5 mM glucose | 8 | 6 | - | - |
| | ICC | 0.5× SC | 0.5 mM Myr-K | 5 mM xylose | 8 | 6 | - | - |
| Fig. 2D | Monogenic | M | None | 1% sucrose | 1 | - | D | 500 |
| Fig. 2E | Monogenic | M | None | 1% sucrose | 5 | - | D | 500 |
| Fig. 2F | Monogenic | M | None | 1% sucrose | 14 | - | D | 500 |
| Fig. 2G | Monogenic | M | None | 1% sucrose | 11 | - | L | 500 |
| Fig. 3A | ICC | SC | 0.5 mM Myr-K (6 weeks) -> 0.5 mM $C_1$-BODIPY 500/510 $C_{12}$ (10 min) | None | 6 | - | C | 200 |
| Fig. 3B | ICC | SC | 0.5 mM Myr-K (3 weeks) -> none (11 days) -> none (3 h) | None | 4 | 5-6 | - | - |
| | ICC | SC | 0.5 mM Myr-K (3 weeks) -> none (11 days) -> 0.5 mM Myr-K (3 h) | None | 4 | 5-6 | - | - |
| Fig. 3C | ICC | 0.5× SC | 0.5 mM neutralized myristic acid | None | 8 | 8 | - | - |
| | ICC | 0.5× SC | 0.5 mM neutralized $[1\text{-}^{13}C_1]$myristic acid | None | 8 | 7 | - | - |
| Fig. 3D | Liquid | Water | None (5 days) -> none (12 h) | None | 5 days | 4 | - | - |
| | Liquid | Water | None (5 days) -> 0.5 mM Myr-K (12 h) | None | 5 days | 4 | - | - |
| | Liquid | Water | None (5 days) -> 50 µM CCCP (12 h) | None | 5 days | 4 | - | - |
| | Liquid | Water | None (5 days) -> 50 µM CCCP + 0.5 mM Myr-K (12 h) | None | 5 days | 4 | - | - |
| Fig. 3E | Solid | SC | 1 mM $[1\text{-}^{13}C_1]$myristic acid | None | 10 | 1 | - | - |
| Fig. 4A, B | ICC | 0.5x SC | 0.1 mM Myr-K | None | 8 | 6 | - | - |
| | ICC | 0.5x SC | 0.1 mM Myr-K + 0.5 mM β-MAG C16:0 | None | 8 | 4 | - | - |
| | ICC | 0.5x SC | 0.1 mM Myr-K + 0.5 mM Pal-K | None | 8 | 6 | - | - |
| | ICC | 0.5x SC | 0.5 mM Myr-K | None | 8 | 6 | - | - |
| | ICC | 0.5x SC | 0.5 mM Myr-K + 0.5 mM β-MAG C16:0 | None | 8 | 4 | - | - |
| | ICC | 0.5x SC | 0.5 mM Myr-K + 0.5 mM Pal-K | None | 8 | 6 | - | - |
| Fig. 4C | ICC | 0.5x SC | 0.1 mM Myr-K | None | 8 | 73 | - | - |
| | ICC | 0.5x SC | 0.1 mM Myr-K + 0.5 mM β-MAG C16:0 | None | 8 | 94 | - | - |
| | ICC | 0.5x SC | 0.1 mM Myr-K + 0.5 mM Pal-K | None | 8 | 175 | - | - |
| | ICC | 0.5x SC | 0.5 mM Myr-K | None | 8 | 172 | - | - |
| | ICC | 0.5x SC | 0.5 mM Myr-K + 0.5 mM β-MAG C16:0 | None | 8 | 109 | - | - |
| | ICC | 0.5x SC | 0.5 mM Myr-K + 0.5 mM Pal-K | None | 8 | 152 | - | - |
| Fig. 4D, E | ICC | 0.5× SC | 0.5 mM neutralized myristic acid | None | 8 | 4 | - | - |

| | | | | | | | | |
|---|---|---|---|---|---|---|---|---|
| | ICC | 0.5× SC | 0.5 mM neutralized $[1\text{-}^{13}C_1]$myristic acid | None | 8 | 4 | - | - |
| | ICC | 0.5× SC | 0.1 mM Myr-K + 0.5 mM neutralized palmitic acid | None | 8 | 4 | - | - |
| | ICC | 0.5× SC | 0.1 mM Myr-K + 0.5 mM neutralized $[1\text{-}^{13}C_1]$ palmitic acid | None | 8 | 3 | - | - |
| Fig. 4F | ICC | 0.5× SC | 0.5 mM neutralized myristic acid + 0.5 mM neutralized palmitic acid | None | 8 | 5 | - | - |
| | ICC | 0.5× SC | 0.5 mM neutralized myristic acid + 0.5 mM neutralized $[1\text{-}^{13}C_1]$palmitic acid | None | 8 | 5 | - | - |
| | ICC | 0.5× SC | 0.5 mM neutralized $[1\text{-}^{13}C_1]$myristic acid + 0.5 mM neutralized palmitic acid | None | 8 | 5 | - | - |

| Figure | Culture | Medium | Fatty acid | Sugar | Week[c] | Replication | Microscopy[d] | Scale (µm) |
|---|---|---|---|---|---|---|---|---|
| Fig. S1A | - | - | None, Parent spore | None | 0 | 8 | - | - |
| | Solid | SC | None, Control | None | 8 | 4 | - | - |
| | Solid | SC | 1 mM pottasium laurate, C12:0-K | None | 8 | 4 | - | - |
| | Solid | SC | 1 mM Myr-K, C14:0-K | None | 8 | 4 | - | - |
| | Solid | SC | 1 mM pottasium palmitate, C16:0-K | None | 8 | 4 | - | - |
| Fig. S1B | Solid | SC | Without glycerol, no fatty acid | None | 8 | 4 | - | - |
| | Solid | SC | With 1 mM glycerol, no fatty acid | None | 8 | 4 | - | - |
| | Solid | SC | Without glycerol, with 1 mM Myr-K | None | 8 | 4 | - | - |
| | Solid | SC | With 1 mM glycerol and 1 mM Myr-K | None | 8 | 3 | - | - |
| Fig. S1C | Solid | SC | None, Control | None | 0, 8 | - | D | 1,000 |
| | Solid | SC | 1 mM pottasium laurate, C12:0-K | None | 0, 8 | - | D | 1,000 |
| | Solid | SC | 1 mM Myr-K, C14:0-K | None | 0, 8 | - | D | 1,000 |
| | Solid | SC | 1 mM pottasium palmitate, C16:0-K | None | 0, 8 | - | D | 1,000 |
| Fig. S1D | Solid | SC | Without glycerol | None | 8 | - | D | 1,000 |
| | Solid | SC | Without glycerol, with 1 mM Myr-K | None | 8 | - | D | 1,000 |
| Fig. S1E | Solid | SC | 1 mM lauric acid, C12:0 | None | 8 | - | D | 1,000 |
| | Solid | SC | 1 mM myristic acid, C14:0 | None | 8 | - | D | 1,000 |
| | Solid | SC | 1 mM palmitic acid, C16:0 | None | 8 | - | D | 1,000 |
| | Solid | SC | 1 mM 12-methyltetradecanonic acid, Antesio-C15:0 | None | 8 | - | D | 1,000 |
| | Solid | SC | 1 mM palmitoleic acid, C16:1Δ9Z | None | 8 | - | D | 1,000 |
| | Solid | SC | 1 mM palmitvaccenic acid, C16:1Δ11Z | None | 8 | - | D | 1,000 |
| Fig. S1F | Solid | SC | 0.05% BSA, BSA | None | 8 | - | D | 1,000 |
| | Solid | SC | 0.5 mM lauric acid-BSA, C12:0-BSA | None | 8 | - | D | 1,000 |
| | Solid | SC | 0.5 mM myristic acid-BSA, C14:0-BSA | None | 8 | - | D | 1,000 |
| | Solid | SC | 0.5 mM palmitic acid-BSA, C16:0-BSA | None | 8 | - | D | 1,000 |
| | Solid | SC | 0.5 mM stearic acid-BSA, C18:0-BSA | None | 8 | - | D | 1,000 |
| | Solid | SC | 0.5 mM oleic acid-BSA, C18:1-BSA | None | 8 | - | D | 1,000 |
| Fig. S1G | Solid | SC | 1 mM C14:0 *sn* -2 monoacylglycerol, β-MAG C14:0 | None | 8 | - | D | 1,000 |
| | Solid | SC | 1 mM C16:0 *sn* -2 monoacylglycerol, β-MAG C16:0 | None | 8 | - | D | 1,000 |
| Fig. S1H | Solid | SC | 0.05% BSA | None | 8 | - | D | 1,000 |
| | Solid | SC | 1 µM lauric acid-BSA | None | 8 | - | D | 1,000 |
| | Solid | SC | 10 µM lauric acid-BSA | None | 8 | - | D | 1,000 |
| | Solid | SC | 100 µM lauric acid-BSA | None | 8 | - | D | 1,000 |
| Fig. S2A | Liquid | SC | None | None | 8 | 3 | - | - |
| | Liquid | SC | 0.5 mM Myr-K | None | 8 | 3 | - | - |
| Fig. S2B | Liquid | SC | None | None | 0, 1, 2, 4, 8 | - | D | 1,000 |
| | Liquid | SC | 0.5 mM Myr-K | None | 0, 1, 2, 4, 8 | - | D | 1,000 |
| Fig. S2C | Liquid | SC | None | None | 8 | - | I | 200 |
| | Liquid | SC | 0.5 mM Myr-K | None | 8 | - | I | 200 |
| Fig. S3A | ICC | 0.5× SC | 0.5 mM Myr-K | None | 0 | - | D | 10,000 |
| Fig. S3B | ICC | 0.5× SC | 0.5 mM Myr-K | None | 8 | - | D | 10,000 |
| Fig. S3C | ICC | 0.5× SC | 0.5 mM Myr-K | None | 8 | - | D | 1,000 |
| Fig. S3D | ICC | 0.5× SC | 0.5 mM Myr-K | None | 8 | - | D | 1,000 |
| Fig. S3E | ICC | 0.5× SC | 0.5 mM Myr-K | None | 8 | - | I | 1,000 |
| Fig. S3F | ICC | 0.5× SC | 0.5 mM Myr-K | None | 8 | - | I | 1,000 |
| Fig. S3G | ICC | 0.5× SC | 0.5 mM Myr-K | None | 8 | - | I | 100 |
| Fig. S3H | ICC | 0.5× SC | 0.5 mM Myr-K | None | 8 | - | I | 1,000 |
| Fig. S3I | ICC | 0.5× SC | 0.5 mM Myr-K | None | 8 | - | I | 100 |
| Fig. S3J | ICC | 0.5× SC | 0.5 mM Myr-K | None | 8 | - | I | 100 |
| Fig. S4A | - | - | None | None | 0 | 5 | - | - |
| | ICC | 0.5× SC | None | None | 12 | 6 | - | - |
| | ICC | 0.5× SC | 0.5 mM Myr-K | None | 12 | 4 | - | - |
| Fig. S4B | ICC | 0.5× SC | None | None | 12 | 6 | - | - |
| | ICC | 0.5× SC | 0.5 mM Myr-K | None | 12 | 4 | - | - |
| Fig. S4C | - | - | None | None | 0 | 26 | - | - |
| | ICC | 0.5× SC | None | None | 12 | 26 | - | - |
| | ICC | 0.5× SC | 0.5 mM Myr-K | None | 12 | 177 | - | - |
| Fig. S4D | ICC | 0.5× SC | None | None | 12 | - | D | 1,000 |
| Fig. S4E | ICC | 0.5× SC | 0.5 mM Myr-K | None | 12 | - | D | 1,000 |
| Fig. S4F | ICC | 0.5× SC | 0.5 mM Myr-K | None | 12 | - | D | 1,000 |
| Fig. S4G | ICC | 0.5× SC | 0.5 mM Myr-K | None | 12 | - | D | 1,000 |
| Fig. S4H | ICC | 0.5× SC | 0.5 mM Myr-K | None | 12 | - | D | 1,000 |
| Fig. S4I | ICC | 0.5× SC | 0.5 mM Myr-K | None | 12 | - | D | 1,000 |
| Fig. S4J | ICC | 0.5× SC | 0.5 mM Myr-K | None | 12 | - | I | 100 |
| Fig. S4K | ICC | 0.5× SC | 0.5 mM Myr-K | None | 12 | - | I | 100 |
| Fig. S5A | ICC | 0.5× SC | None | None | 12 | - | F | 1,000 |
| Fig. S5B | ICC | 0.5× SC | None | None | 12 | - | IK | 1,000 |
| Fig. S5C | ICC | 0.5× SC | None | None | 12 | - | F | 200 |
| Fig. S5D | ICC | 0.5× SC | 0.5 mM Myr-K | None | 12 | - | IK | 1,000 |
| Fig. S5E | ICC | 0.5× SC | 0.5 mM Myr-K | None | 12 | - | F | 1,000 |
| Fig. S5F | ICC | 0.5× SC | 0.5 mM Myr-K | None | 12 | - | IK | 200 |
| Fig. S5G | ICC | 0.5× SC | 0.5 mM Myr-K | None | 12 | - | IK | 200 |
| Fig. S5H | ICC | 0.5× SC | 0.5 mM Myr-K | None | 12 | - | IK | 50 |
| Fig. S6A | ICC | 0.5× SC | None | None | 8 | - | D | 1,000 |
| | ICC | 0.5× SC | 0.5 mM Myr-K | None | 8 | - | D | 1,000 |
| | ICC | 0.5× SC | None | 5 mM glucose | 8 | - | D | 1,000 |

| | | | | | | | | |
|---|---|---|---|---|---|---|---|---|
| | ICC | 0.5× SC | 0.5 mM Myr-K | 5 mM glucose | 8 | - | D | 1,000 |
| | ICC | 0.5× SC | None | 5 mM xylose | 8 | - | D | 1,000 |
| | ICC | 0.5× SC | 0.5 mM Myr-K | 5 mM xylose | 8 | - | D | 1,000 |
| Fig. S6B | ICC | 0.5× SC | 0.5 mM Myr-K | None | 8 | 211 | - | - |
| | ICC | 0.5× SC | 0.5 mM Myr-K | 5 mM glucose | 8 | 218 | - | - |
| | ICC | 0.5× SC | 0.5 mM Myr-K | 5 mM xylose | 8 | 188 | - | - |
| Fig. S7A | - | - | None | None | 0 | - | L, E | 20 |
| | ICC | 0.5x SC | 0.5 mM Myr-K | None | 12 | - | L, E | 20 |
| | ICC | 0.5x SC | 0.1 mM Myr-K + 0.5 mM Pal-K | None | 16 | - | L, E | 20 |
| Fig. S7B | - | - | None | None | 0 | - | T | 1 |
| | ICC | 0.5x SC | 0.5 mM Myr-K | None | 19 | - | T | 1 |
| | ICC | 0.5x SC | 0.1 mM Myr-K + 0.5 mM Pal-K | None | 16 | - | T | 1 |
| Fig. S8A | ICC | SC | 0.5 mM Myr-K (6 weeks) -> 0.5 mM $C_1$-BODIPY 500/510 $C_{12}$ (1 days) | None | 6 | - | E | 100 |
| Fig. S8B | ICC | SC | 0.5 mM Myr-K (6 weeks) -> 0.5 mM $C_1$-BODIPY 500/510 $C_{12}$ (5 days) | None | 6 | - | E | 10 |
| Fig. S8C | ICC | SC | 0.5 mM Myr-K (6 weeks) -> 0.5 mM $C_1$-BODIPY 500/510 $C_{12}$ (1 days) | None | 6 | - | E | 100 |
| Fig. S8D | ICC | SC | 0.5 mM Myr-K (6 weeks) -> 0.5 mM $C_1$-BODIPY 500/510 $C_{12}$ (1 days) | None | 6 | - | E | 100 |
| Fig. S8E | ICC | SC | 0.5 mM Myr-K (6 weeks) -> 0.5 mM $C_1$-BODIPY 500/510 $C_{12}$ (5 days) | None | 6 | - | E | 10 |
| Fig. S8F | Liquid | Water | Water (1 week) -> 0.5 mM $C_1$-BODIPY 500/510 $C_{12}$ (10 min) | None | 1 | - | E | 100 |
| Fig. S8G | Liquid | Water | Water (1 week) -> 0.5 mM $C_1$-BODIPY 500/510 $C_{12}$ (7 days) | None | 1 | - | E | 100 |
| Fig. S9A | ICC | SC | 0.5 mM Myr-K (6 weeks) -> 0.5 mM BODIPY FL $C_{16}$ (10 min) | None | 6 | - | E | 100 |
| Fig. S9B | ICC | SC | 0.5 mM Myr-K (7 weeks) -> 0.5 mM BODIPY FL $C_{16}$ (4 h) | None | 7 | - | E | 100 |
| Fig. S9C | ICC | SC | 0.5 mM Myr-K (7 weeks) -> 0.5 mM BODIPY FL $C_{16}$ (4 h) | None | 7 | - | E | 20 |
| Fig. S9D | ICC | SC | 0.5 mM Myr-K (8 weeks) -> 0.5 mM BODIPY FL $C_{16}$ (3 days) | None | 8 | - | E | 100 |
| Fig. S9E | ICC | SC | 0.5 mM Myr-K (8 weeks) -> 0.5 mM BODIPY FL $C_{16}$ (3 days) | None | 8 | - | E | 20 |
| Fig. S9F | ICC | SC | 0.5 mM Myr-K (8 weeks) -> 0.5 mM BODIPY FL $C_{16}$ (3 days) | None | 8 | - | E | 100 |
| Fig. S9G | ICC | SC | 0.5 mM Myr-K (6 weeks) -> 0.5 mM BODIPY FL $C_{16}$ (9 days) | None | 7 | - | E | 100 |
| Fig. S9H | ICC | SC | 0.5 mM Myr-K (6 weeks) -> 0.5 mM BODIPY FL $C_{16}$ (9 days) | None | 7 | - | E | 20 |
| Fig. S9I | Liquid | Water | Water (1 week) -> 0.5 mM BODIPY FL $C_{16}$ (10 min) | None | 1 | - | E | 100 |
| Fig. S9J | Liquid | Water | Water (1 week) -> 0.5 mM BODIPY FL $C_{16}$ (7 days) | None | 2 | - | E | 100 |
| Fig. S10A | ICC | 0.5× SC | 0.1 mM Myr-K | None | 8 | - | F | 200 |
| Fig. S10B | ICC | 0.5× SC | 0.1 mM Myr-K | None | 8 | - | F | 200 |
| Fig. S10C | ICC | 0.5× SC | 0.5 mM Myr-K | None | 8 | - | F | 200 |
| Fig. S10D | ICC | 0.5× SC | 0.5 mM Myr-K | None | 8 | - | F | 200 |
| Fig. S10E | ICC | 0.5× SC | 0.1 mM Myr-K + 0.5 mM β-MAG C16:0 | None | 8 | - | F | 200 |
| Fig. S10F | ICC | 0.5× SC | 0.1 mM Myr-K + 0.5 mM β-MAG C16:0 | None | 8 | - | F | 200 |
| Fig. S10G | ICC | 0.5× SC | 0.1 mM Myr-K + 0.5 mM β-MAG C16:0 | None | 8 | - | F | 200 |
| Fig. S10H | ICC | 0.5× SC | 0.1 mM Myr-K + 0.5 mM β-MAG C16:0 | None | 8 | - | F | 200 |
| Fig. S10I | ICC | 0.5× SC | 0.1 mM Myr-K + 0.5 mM Pal-K | None | 8 | - | F | 200 |
| Fig. S10J | ICC | 0.5× SC | 0.1 mM Myr-K + 0.5 mM Pal-K | None | 8 | - | F | 200 |
| Fig. S10K | ICC | 0.5× SC | 0.1 mM Myr-K + 0.5 mM Pal-K | None | 8 | - | F | 200 |
| Fig. S10L | ICC | 0.5× SC | 0.1 mM Myr-K + 0.5 mM Pal-K | None | 8 | - | F | 200 |
| Fig. S10M | ICC | 0.5× SC | 0.5 mM Myr-K + 0.5 mM β-MAG C16:0 | None | 8 | - | F | 200 |
| Fig. S10N | ICC | 0.5× SC | 0.5 mM Myr-K + 0.5 mM β-MAG C16:0 | None | 8 | - | F | 200 |
| Fig. S10O | ICC | 0.5× SC | 0.5 mM Myr-K + 0.5 mM β-MAG C16:0 | None | 8 | - | F | 200 |
| Fig. S10P | ICC | 0.5× SC | 0.5 mM Myr-K + 0.5 mM β-MAG C16:0 | None | 8 | - | F | 200 |
| Fig. S10Q | ICC | 0.5× SC | 0.5 mM Myr-K + 0.5 mM Pal-K | None | 8 | - | F | 200 |
| Fig. S10R | ICC | 0.5× SC | 0.5 mM Myr-K + 0.5 mM Pal-K | None | 8 | - | F | 200 |
| Fig. S10S | ICC | 0.5× SC | 0.5 mM Myr-K + 0.5 mM Pal-K | None | 8 | - | F | 200 |
| Fig. S10T | ICC | 0.5× SC | 0.5 mM Myr-K + 0.5 mM Pal-K | None | 8 | - | F | 200 |
| Movie S1 | ICC | SC | 0.5 mM Myr-K (6 weeks) -> 0.5 mM $C_1$-BODIPY 500/510 $C_{12}$ (10 min) | None | 6 | - | C | 25 |

**Table S3.** Infection capability of myristate-induced spores. Carrot hairy roots were inoculated with a single myristate-induced spore and grown for two months. Six independent experiments were performed.

| Trial | Number of inoculated spores | Germination rate (%) | Rate of spores which colonize to roots and produce next generation spores (%) | |
|---|---|---|---|---|
| | | | per inoculated spore | per germinated spore |
| Trial 1 | 20 | 30 | 15 | 50 |
| Trial 2 | 40 | 33 | 18 | 54 |
| Trial 3 | 29 | 7 | 3 | 50 |
| Trial 4 | 40 | 20 | 13 | 63 |
| Trial 5 | 60 | 12 | 2 | 14 |
| Trial 6 | 56 | 5 | 4 | 67 |

**Table S4.** Primers used in this study.

| Gene | RIR ID[a] | Annotation | Sequence | Reference |
|---|---|---|---|---|
| *FAD1* | RIR_0930900 | Acyl-CoA dehydrogenase | TGGTATTCATTCGTGCCATGA | This study |
| | | | GCAGCGCGTGCAAGTG | |
| *FAL3* | RIR_0764900 | Acyl-CoA ligase | TCAAAAAGATGCCAAAGTGACAAT | This study |
| | | | AACCTTGTGTTACGAGCATCCA | |
| *FAL4* | RIR_2539500 | Acyl-CoA ligase | ATTGTAGAGGGTTATGGGCAGACT | This study |
| | | | TTTCGCCTCGAAGACCAACT | |
| *ICL1* | RIR_1924000 | Isocitrate lyase | CCGCGCCATAGCGTATG | This study |
| | | | TAGGTTTTGCAGTCTCCATCCA | |
| *MS1* | RIR_2183100 | Malate synthase | GAAGCGAGACTTTGGAATGACAT | This study |
| | | | ACCACGAGGAAGGCCAATAA | |
| *PCK2* | RIR_2338300 | Phosphoenolpyruvate carboxykinase | AAAAACTTGGAAAGGTCCTCAAGA | This study |
| | | | AGAAATCTCGCAGCTAACGAATCT | |
| *FBP1* | RIR_2783400 | Fructose-1,6-bisphosphate phosphatase | CGCGTATCCCGCTGATAAAA | This study |
| | | | GGAAAGCCATAGGAAAACATTCA | |
| *CIT1* | RIR_0822400 | Citrate synthase | CTACATGGCCTTGCTAATCAAGAA | This study |
| | | | CACCAATCGCGTCTCTCATCT | |
| *ACH1* | RIR_2973500 | Aconitate hydratase | CTTCCCGGGTGGTCTTATGA | This study |
| | | | ACCCAAACCACCAGCGTTAG | |
| *ICD2* | RIR_1281200 | Isocitrate dehydrogenase | AGCGGATCGGCCAGAAC | This study |
| | | | CAAGGGTAAAATCGGTTGTTGTAGA | |
| *KGD2* | RIR_2870300 | 2-Ketoglutarate dehydrogenase | TTGTACGGTGATCGCGAAGA | This study |
| | | | GGCAATACGTAGACGCATCCTAT | |
| *SDH2* | RIR_2582900 | Succinate dehydrogenase | GCAGGATGCTACATTGACTTTCC | This study |
| | | | AGCGCAAGATCCGCAAATA | |
| *MDH1* | RIR_0715300 | Malate dehydrogenese | GAAGTTGTTAAAGCCAAGGATGGT | This study |
| | | | ACCAGCCTGTGCCATTGAA | |
| *NMT1* | RIR_3261900 | *N* -myristoyltransferase | CGATGCGATGTTTCGCTTT | This study |
| | | | GGTGGTTGAAGAGCCCATTTT | |
| *EF-1β* | RIR_3067400 | Translation elongation factor 1 subunit beta | CCCATGCAGCTCGATGGTA | Kobae et al., 2015[b] |
| | | | TGCCAGGAAGTGAAGAAAATGA | |
| *Act* | RIR_2118800 | Actin | TGACAACGGTTCCGGTATGTG | Tsuzuki et al., 2016[c] |
| | | | ATCTTTCTGACCCATCCCAACC | |

[a]Maeda et al., Evidence of non-tandemly repeated rDNAs and their intragenomic heterogeneity in *Rhizophagus irregularis. Commun. Biol.* **1**, 87 (2018).

[b]Kobae et al., Up-regulation of genes involved in *N* -acetylglucosamine uptake and metabolism suggests a recycling mode of chitin in intraradical mycelium of arbuscular mycorrhizal fungi. *Mycorrhiza* **25**, 411-417 (2015).

[c]Tsuzuki et al., Strigolactone-induced putative secreted protein 1 is required for the establishment of symbiosis by the arbuscular mycorrhizal fungus *Rhizophagus irregularis* . *Molecular Plant-Microbe Interactions* **29**, 277-286 (2016).

**Movie S1.** Translocation of the fluorescently labelled fatty acid derivative $C_1$-BODIPY 500/510 $C_{12}$ taken up by *R. irregularis*. Fluorescent images of $C_1$-BODIPY 500/510 $C_{12}$ were superimposed bright field images of the runner hyphae. AM fungi were incubated using an immobilized cell culture system with the modified SC medium containing 0.5 mM fluorescent probe.
